# Targeting complement C3a receptor resolves mitochondrial hyperfusion and subretinal microglial activation in progranulin-deficient frontotemporal dementia

**DOI:** 10.1101/2024.05.29.595206

**Authors:** Li Xuan Tan, Frederike Cosima Oertel, An Cheng, Yann Cobigo, Azeen Keihani, Daniel J Bennett, Ahmed Abdelhak, Shivany Condor Montes, Makenna Chapman, Robert Y Chen, Christian Cordano, Michael E. Ward, Kaitlin Casaletto, Joel H. Kramer, Howard J Rosen, Adam Boxer, Bruce L Miller, Ari J Green, Fanny M Elahi, Aparna Lakkaraju

## Abstract

Mutations in progranulin (*GRN*) cause frontotemporal dementia (*GRN*-FTD) due to deficiency of the pleiotropic protein progranulin. *GRN*-FTD exhibits diverse pathologies including lysosome dysfunction, lipofuscinosis, microgliosis, and neuroinflammation. Yet, how progranulin loss causes disease remains unresolved. Here, we report that non-invasive retinal imaging of *GRN*-FTD patients revealed deficits in photoreceptors and the retinal pigment epithelium (RPE) that correlate with cognitive decline. Likewise, *Grn^−/−^* mice exhibit early RPE dysfunction, microglial activation, and subsequent photoreceptor loss. Super-resolution live imaging and transcriptomic analyses identified RPE mitochondria as an early driver of retinal dysfunction. Loss of mitochondrial fission protein 1 (MTFP1) in *Grn^−/−^* RPE causes mitochondrial hyperfusion and bioenergetic defects, leading to NF-kB-mediated activation of complement C3a-C3a receptor signaling, which drives further mitochondrial hyperfusion and retinal inflammation. C3aR antagonism restores RPE mitochondrial integrity and limits subretinal microglial activation. Our study identifies a previously unrecognized mechanism by which progranulin modulates mitochondrial integrity and complement-mediated neuroinflammation.

## INTRODUCTION

Frontotemporal dementia (FTD), the second most prevalent form of early-onset dementia, is associated with the progressive impairment of behavioral, language, and executive functions ^1,2^. Heterozygous mutations in the progranulin gene (*GRN*) account for ∼5-20% of FTD cases (*GRN*-FTD), whereas homozygous *GRN* mutations cause neuronal ceroid lipofuscinosis type 11 (CLN11), which is characterized by retinal degeneration, ataxia, and cognitive deficits in early adulthood ^3,4^. Using non-invasive optical coherence tomography (OCT), we have previously identified structural abnormalities in the retina including loss of total macular volume and thinning of the retinal nerve fiber layer in patients with *GRN*-FTD. These retinal phenotypes are also observed in pre-symptomatic carriers, suggesting that the retina is uniquely sensitive to progranulin insufficiency ^5^.

Progranulin, a cysteine-rich glycoprotein, and the granulin peptides generated by its proteolysis, have been implicated in pleiotropic tissue-specific functions including signaling, neuroprotection, and inflammation ^6,7^. Phenotypic and multi-omics analyses of *GRN*-FTD patient tissues, *Grn^−/−^* mice, and cell-based models suggest that progranulin loss impairs lysosome function, autophagy, and lipid catabolism, leads to the accumulation of lysosomal storage material (lipofuscinosis), and promotes inflammation mediated by reactive microglia ^8–13^. Upregulation of the complement pathway, microglial activation, synaptic pruning, Nf-kB signaling, and hyperexcitability have been reported in the thalamus and striatum of *Grn^−/−^* mice ^14,15^. However, precisely how progranulin modulates microglial homeostasis and the immune response, and how these are compromised in *GRN*-FTD are not well understood.

In this study, we addressed these questions by investigating the mechanisms by which progranulin loss leads to microglial activation and neurodegeneration in the retina. As the most accessible part of the CNS, the retina is uniquely amenable to non-invasive imaging that affords a powerful opportunity to monitor neurodegenerative changes longitudinally in living humans. We performed high-resolution retinal imaging of *GRN* mutation carriers to establish the extent of retinal deficits and conducted multimodal analyses of *Grn^−/−^* mice to molecularly dissect the link between progranulin, microglia, and retinal degeneration. In the healthy retina, microglia reside in the inner and outer plexiform layers (IPL and OPL, Figure 1A), where they participate in immune surveillance to maintain retinal homeostasis. Migration of microglia from the IPL and OPL into the subretinal space between the photoreceptors and the retinal pigment epithelium (RPE) is associated with photoreceptor loss and visual deficits ^16,17^. The RPE is a postmitotic, highly metabolically active tissue that performs numerous functions indispensable for photoreceptor health and vision ^18,19^. In humans with *GRN* mutations, semi-automated OCT segmentation revealed thinning of the RPE and photoreceptor layers and thickening of the OPL, suggesting RPE and photoreceptor degeneration and microglial expansion. These retinal deficits correlated with cognitive decline and white matter changes in the brain.

**Figure1.**
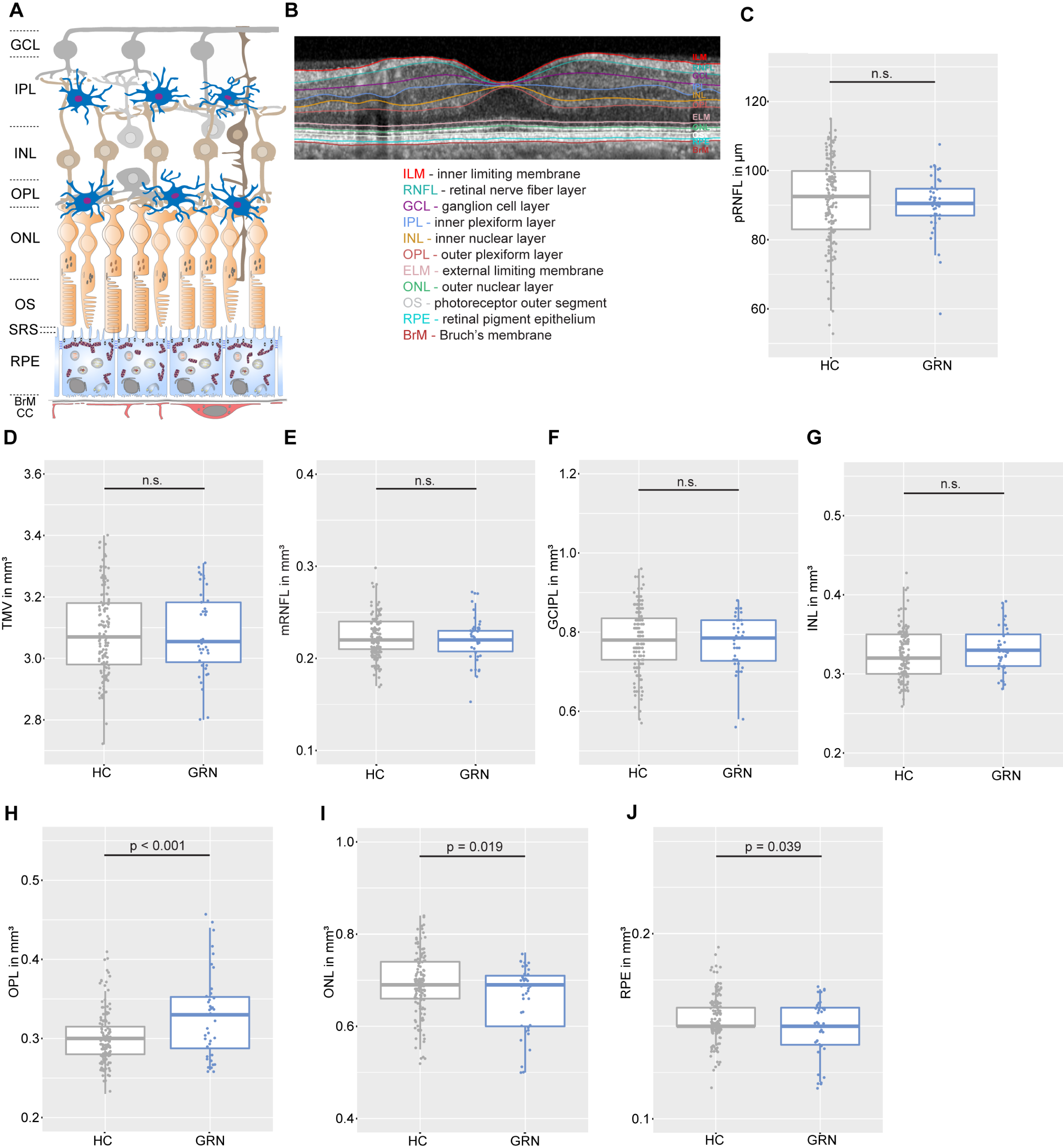
Outer retinal deficits in *GRN* mutation carriers. (**A**) Schematic of layers of the neural retina and RPE. (**B**) Representative OCT scan of a healthy control with retinal layers. Boxplots of cross-sectional OCT data in healthy controls (HCs) and heterozygous *GRN* mutation carriers (*GRN*) for (**C**) peripapillary retinal nerve fiber layer thickness (pRNFL); (**D**) total macular volume (TMV); (**E**) macular retinal nerve fiber layer volume (mRNFL); (**F**) combined ganglion cell and inner plexiform layer volume (GCIPL); (**G**) inner nuclear layer volume (INL); (**H**) outer plexiform layer volume (OPL); (**I**) outer nuclear layer volume (ONL); and (**J**) retinal pigment epithelium volume (RPE). Values of individual eyes overlayed as jitter. The data are presented as the Mean ± SEM. *p* values were calculated using Student’s *t*-test.

The retinal phenotype of *Grn^−/−^* mice recapitulates several features of *GRN*-FTD and CLN11 including lipofuscinosis, microglial migration, retinal degeneration, and visual function deficits ^5,20–22^. Using advanced live imaging of RPE explants, RNAseq analyses, and biochemical approaches, our studies identified mitochondrial dysfunction in the RPE as a central and early driver of microglial activation and photoreceptor degeneration in *Grn^−/−^* mice. Loss of progranulin disrupted mTOR-mediated translation of mitochondrial fission process 1 (MTFP1), resulting in mitochondrial hyperfusion and bioenergetic deficits in 4-month-old *Grn^−/−^* mice RPE. Mitochondrial stress activated the transcription of NF-κ;B target genes including the complement protein C3, the core effector of the complement pathway, and its cognate receptor C3aR. Abnormal C3a-C3aR signaling reinforced a feed-forward cycle of further mitochondrial hyperfusion and activation of subretinal microglia. These microglia phagocytosed photoreceptor outer segments and RPE microvilli, thus initiating retinal degeneration. Blocking C3aR activation restored RPE mitochondrial function and microglial homeostasis in *Grn^−/−^*mice.

These studies reveal a previously unrecognized function of progranulin in maintaining mitochondrial integrity, establish an intricate molecular pathway to explain how progranulin deficiency leads to complement-mediated microglial activation and neurodegeneration, and identify C3aR as a potential therapeutic target in *GRN*-FTD.

## RESULTS

### Photoreceptor loss and RPE thinning in human heterozygous *GRN* mutation carriers

To evaluate the extent of retinal deficits in patients with *GRN*-FTD, we conducted an OCT study on 46 eyes from 23 heterozygous *GRN* mutation carriers and 147 eyes from 85 healthy controls (Table S1). Initially, nine patients (39%) had a clinical dementia rating (CDR) score of 0, with a median CDR score of 0.5. Longitudinal OCT data were available for nine heterozygous *GRN* mutation carriers (18 eyes) with a median follow-up of 41 months. Nine patients were previously included in a cross-sectional study by Ward et al ^13^. We first analyzed qualitative OCT changes: eight eyes (17%) of five subjects (22%) exhibited drusen-like deposits, as described by Ward et al.^5^. Five eyes (11%) showed other pathologies, with the most common being epiretinal membranes in three eyes.

Analyses of retinal layer thickness between heterozygous *GRN* mutation carriers and healthy controls (Figure 1B) at baseline showed that the total macular volume (TMV), the peripapillary retinal nerve fiber layer (pRNFL), and the combined ganglion cell and inner plexiform layers (GCIP), which house the axons and cell bodies of retinal ganglion cells, did not differ between heterozygous *GRN* mutation carriers and healthy controls (Figure 1C-J, Table S2). In contrast, compared to healthy controls, heterozygous *GRN* mutation carriers exhibited thinning of the RPE layer and the outer nuclear layer (ONL), which has photoreceptor cell bodies, indicative of RPE stress and photoreceptor loss. GRN mutation carriers also had thicker outer plexiform layers (OPL), suggesting expansion and/or migration of microglia.

After excluding eyes with drusen-like pathologies, the OPL remained thicker in *GRN* mutation carriers (0.33 ± 0.05 mm^3^) compared to healthy controls (B = 0.029, SE = 0.009, p = 0.002), and the ONL and RPE remained thinner in *GRN* mutation carriers (ONL: 0.66 ± 0.07 mm^3^; RPE: 0.15 ± 0.02 mm^3^) compared to healthy controls (ONL: B = −0.039, SE = 0.018, p = 0.030; RPE: B = −0.009, SE = 0.003, p = 0.009). The OPL/ONL-ratio was significantly higher in *GRN* mutation carriers (0.51 ± 0.13) than in healthy controls (0.44 ± 0.08, B = 0.079, SE = 0.021, p < 0.001).

### Association of outer retinal layer changes with cognition

We investigated the clinical significance of the higher OPL/ONL-ratio in heterozygous *GRN* mutation carriers. At baseline, cognitive parameters were comparable between heterozygous *GRN* mutation carriers with high versus low OPL/ONL-ratio (Table S3), consistent with the preclinical disease stage of the *GRN* cohort. Longitudinally, the heterozygous *GRN* mutation carriers with high versus low OPL/ONL-ratio had greater decline in executive functions (p = 0.039). Detailed analyses of executive functions and language fluency, which are described to be early affected domains specifically in *GRN*-FTLD ^23^, revealed a decline in lexical fluency, semantic fluency, and face affect naming. Of note, visual stimuli are not required during lexical fluency and semantic fluency tasks (Table S4).

### Association of outer retinal layer changes with brain atrophy and white matter disease

Heterozygous *GRN* mutation carriers develop preclinical asymmetric atrophy in the insula, temporal and parietal lobes ^2,24^. In our cohorts, atrophy in these regions of interest (ROIs) did not differ between heterozygous *GRN* mutation carriers with high vs. low OPL/ONL-ratios (Table S5). Eighteen heterozygous *GRN* mutation carriers were evaluated for white matter hyperintensities according to the Fazekas scale and compared with 56 healthy controls at comparable ages (*GRN* carriers: 63 ± 11 years, HC: 62 ± 15 years) ^25^. Severe periventricular white matter hyperintensities (PVWMH) were seen in two patients (Fazekas 3, 11%) and severe deep white matter hyperintensities (DWMH) were seen in five patients (Fazekas 3, 28%). In contrast, severe white matter disease burden (Fazekas 3) was only seen in five healthy controls (9%) for DWMH. All other patients had mild to moderate PVWMH and DWMH. These high frequencies of white matter abnormalities are in line with previously published cohorts ^26^.

*GRN* mutation carriers with a PVWMH score of 3 had a higher OPL/ONL-ratio (mean = 0.84) than patients with a lower PVWMH score (mean = 0.48; t = 11.49, 95% CI [0.288, 0.422], p < 0.001). *GRN* mutation carriers with a DWMH score of 3 also had a numerically higher OPL/ONL-ratio (mean = 0.76) than patients with a lower DWMH score (mean = 0.47); a difference that showed a trend but did not reach statistical significance (t = 3.269, 95% CI [0.758, 0.474], p = 0.055). Taken together, these clinical results suggest that in patients with *GRN*-FTD, progranulin insufficiency is associated with outer retinal thinning and that this correlates with cognitive and white matter deficits.

### Early recruitment of microglia to the subretinal space precedes photoreceptor loss in *Grn^−/−^* **mice**

To identify mechanisms that could underlie the clinical observations in *GRN*-FTD patients, we performed morphometric analyses of retinal layers in *Grn^−/−^* mice. We observed significant loss of ONL and INL thicknesses in 10-month-old *Grn^−/−^* mice compared to age-matched controls (Figures 2A and 2B), in agreement with previous reports ^20,21^.

**Figure 2.**
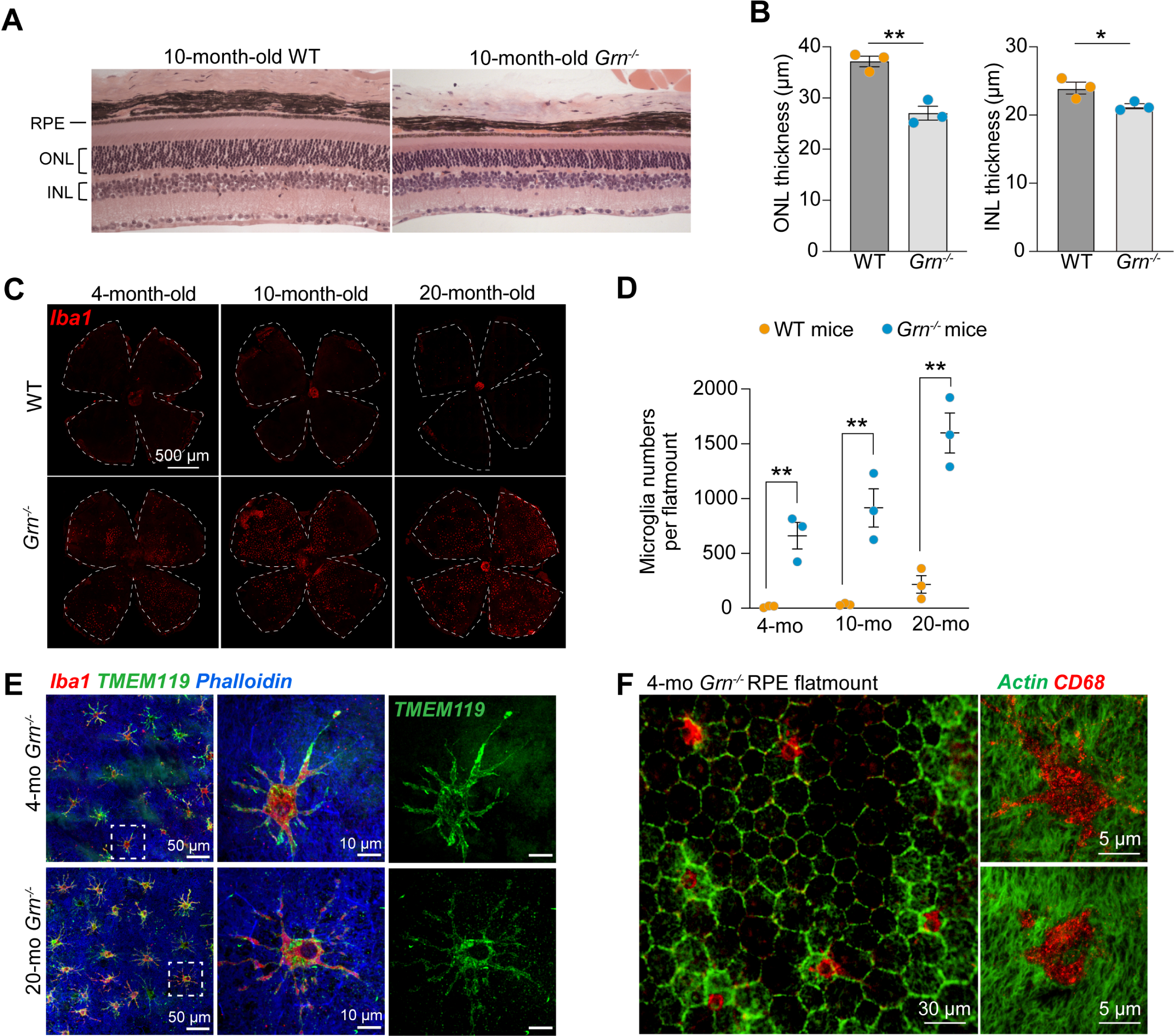
Microglial recruitment to the subretinal space precedes photoreceptor loss in *Grn^−/−^*mice. (**A**) Representative cross-sections of glycol methacrylate-embedded retinas from 10-month-old wildtype (WT) and *Grn^−/−^* mice. (**B**) Quantification of outer (ONL) and inner (INL) nuclear layer thicknesses from (A). Mean ± SEM, 3 mice per genotype. *, *p* < 0.05; **, *p* < 0.01, *t*-test. (**C**) Low magnification images showing representative full RPE flatmounts from 4-, 10-, and 20-month-old WT and *Grn^−/−^* mice immunostained for Iba1 (red). (**D**) Total number of Iba1-positive cells (subretinal microglia) per RPE flatmount in 4-, 10-, and 20-mo WT and *Grn^−/−^*mice. Mean ± SEM, 3 mice/genotype. **, *p* < 0.01 by two-way ANOVA with Tukey’s multiple comparisons test. (**E**) Representative images showing colocalization of Iba1 (red) and TMEM119 (green) in 4- and 20-mo *Grn^−/−^* mice RPE flatmounts. RPE layer is labeled with the actin stain phalloidin (blue). Also see supplementary figure S1. (**F**) CD68 (red) immunostaining of sub-retinal microglia in a 4-month-old *Grn^−/−^* mouse RPE flatmount. Actin stain phalloidin (green) outlines RPE cell boundaries.

As infiltration of microglia from the IPL and OPL to the subretinal space is associated with photoreceptor degeneration ^16,17^, we next investigated whether the time course of photoreceptor degeneration correlated with microglial migration. Immunostaining RPE flatmounts and retinal cryosections from young (3-4 months), 10-mo, and aged (17-20 months) mice showed significant numbers of subretinal microglia in *Grn*^−/−^ mice as early as 3 months, which increased with age and preceded photoreceptor degeneration (Figures 2C, 2D, and S1A). High-magnification imaging of RPE flatmounts confirmed that these microglia had a radial morphology and expressed canonical microglia markers (TMEM119) and markers for activated, highly phagocytic microglia (CD68) (Figures 2E and 2F). These data suggest that invasion of microglia from the IPL and OPL into the subretinal space is an early event that leads to subsequent photoreceptor loss in *Grn*^−/−^ mice.

### Subretinal microglia, and not the RPE, are the source of lipofuscin autofluorescence in *Grn^−/−^* mice retinas

Although autofluorescent lipofuscin-like storage material has been observed in the retinas of *GRN*-FTD patients and *Grn*^−/−^ mice, the origin and cellular location of these deposits has not been established ^13,20^. Circadian phagocytosis and digestion of photoreceptor outer segment tips by the retinal pigment epithelium is essential for maintaining photoreceptor health and function. Overactive phagocytosis or inefficient lysosomal clearance of outer segments leads to the accumulation of vitamin A metabolites in the form of lipofuscin bisretinoids in the RPE, which are associated with blinding diseases called macular degenerations ^27^.

Because progranulin has been reported to regulate the activity of cathepsin D ^28,29^, the major protease responsible for degrading outer segment proteins ^19^, we asked whether impaired phagolysosomal clearance of outer segments could be the source of lipofuscin in *Grn*^−/−^ mice retinas. We quantified outer segment phagosomes by staining RPE flatmounts for rhodopsin and found no significant differences in the number of phagosomes between wildtype and *Grn*^−/−^ RPE from 4- and 15-month-old mice (Figures 3A and 3B). *Grn*^−/−^ RPE had similar levels of cathepsin D protein and normal lysosome morphology compared to age-matched wildtype mice RPE (Figures 3C and 3D), suggesting that RPE lysosome dysfunction is unlikely to be the source of lipofuscin in *GRN*-FTD patients and *Grn*^−/−^ mice.

**Figure 3.**
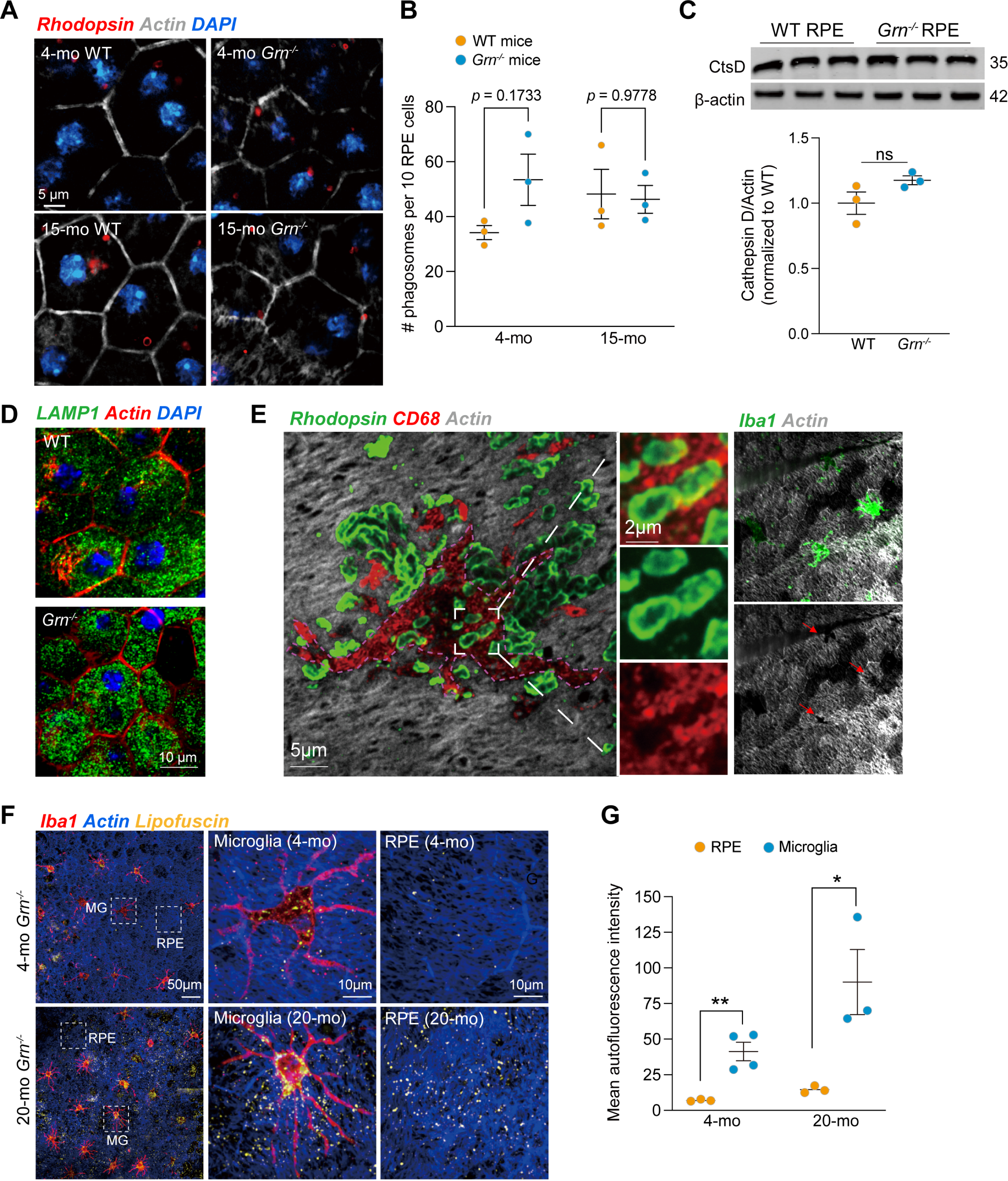
Efficient phagolysosomal clearance in the RPE with lipofuscin accumulation in subretinal microglia in *Grn^−/−^* retinas. (**A**) 4- and 15-mo WT and *Grn^−/−^* mice RPE flatmounts immunostained for photoreceptor outer segment phagosomes (rhodopsin, red). Phalloidin: gray, DAPI: blue. (**B**) Quantification of phagosome numbers per 10 RPE cells at 3 hours after light onset. Mean ± SEM, 3 mice/genotype at each age. **, *p* < 0.01, two-way ANOVA with Tukey’s multiple comparisons test. (**C**) Immunoblotting and quantification of cathepsin D levels in RPE lysates from WT and *Grn^−/−^* mice. Mean ± SEM, n = 3. *, *p* < 0.05, t-test. (**D**) Representative images of LAMP1 immunostaining (green) in WT and *Grn^−/−^* mice RPE flatmounts. (**E**) Representative high-magnification image of 4-mo *Grn^−/−^*mouse RPE flatmount immunostained for CD68 (red) and photoreceptor outer segments (rhodopsin, green). RPE microvilli are stained with phalloidin (gray). High magnification images show phagocytosed outer segments and RPE microvilli within subretinal microglia. Arrows in rightmost images show CD68+ microglia overlying areas of “bald” RPE (red arrows). (**F**) Representative images of Iba1 immunostaining (red) in 4- and 20-mo *Grn^−/−^* mice RPE flatmounts. RPE microvilli are stained by the actin stain phalloidin (blue). Autofluorescence due to lipofuscin accumulation is shown in yellow. Images on the right are high-magnification views of the microglia or RPE in white dashed squares. (**G**) Quantification of autofluorescence signal in RPE and microglia. Mean ± SEM, 3 mice/genotype and age. *, *p* < 0.05, **, *p* < 0.01, two-way ANOVA with Tukey’s multiple comparisons test. Also see Supplementary Figure S1.

High-resolution immunofluorescence imaging of 4-month-old *Grn*^−/−^ RPE flatmounts revealed that subretinal microglia were positive for CD68, indicating that they were actively phagocytic (Figure 3E). In support of this, high-magnification images showed photoreceptor outer segments and RPE microvilli within these microglia. The disruption of the RPE monolayer could also be visualized by hollow spaces in the shape of the infiltrating microglia at the apical surface of the RPE (Figure 3E, red arrows in rightmost panels). Autofluorescence imaging of RPE flatmounts showed that these subretinal microglia, and not the RPE, were the predominant source of lipofuscin autofluorescence in these mice, and this increased with age (Figures 3F and 3G).

### Transcriptomic analyses identify mitochondrial dysfunction and complement activation as early pathogenic mechanisms in *Grn*^−/−^ mice retinas

To identify putative mechanisms by which progranulin loss drives microglial infiltration and retinal neurodegeneration, we conducted a transcriptomic analysis of the RPE in WT and *Grn*^−/−^ mice. We selected 4- and 14-month-old mice, aiming to capture changes before and after the onset of retinal degeneration. The t-SNE map indicated distinctions in gene expression between WT and *Grn*^−/−^ mice starting at 4 months, with clear differences emerging by 14 months (Figure 4A). RNAseq analyses revealed a significant upregulation of genes associated with reactive oxygen species (ROS) metabolic processes, a subset of phagosome and lysosome genes, and genes involved in complement and coagulation cascades in 4-month-old *Grn*^−/−^ RPE (Figure 4B), indicating that mitochondrial dysfunction, oxidative stress, and early activation of the complement pathway drive pathology in *Grn*^−/−^ mice retinas.

**Figure 4.**
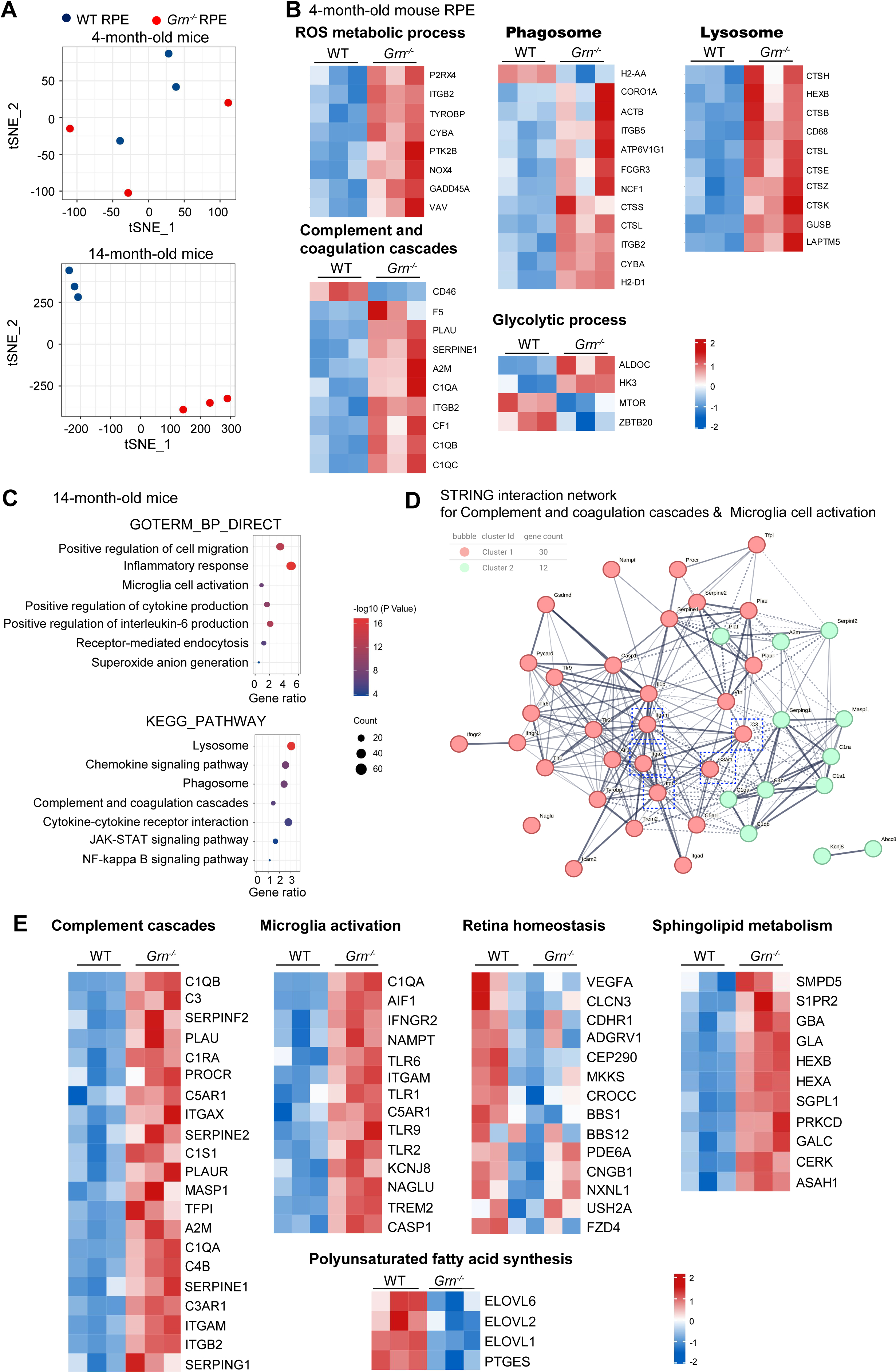
Transcriptomic analyses identify RPE mitochondrial stress as a potential early driver of retinal neurodegeneration in *Grn^−/−^* mice. (**A**) Position and phenotype of clusters on the t-SNE map for 4- and 14-month-old wildtype (WT) and *Grn^−/−^* mice. (**B**) Heatmaps showing alterations in the expression of pathway-specific genes related to reactive oxygen species (ROS) metabolic process, phagosome, lysosome, and complement and coagulation cascades in 4-month-old *Grn^−/−^* RPE. (**C**) GO gene set enrichment analysis results shown as a bubble plot where the x-axis represents gene ratio, and color indicates the adjusted *p*-value for GO terms (left). (**D**) STRING interaction network between complement and coagulation cascades and microglia cell activation clusters is shown as a directed network graph. Nodes represent proteins, and lines indicate interactions in this 1st order interactome. Only those proteins reported at a confidence level of 95% are considered. (**E**) Heatmaps of pathway-specific expression of genes that regulate complement and coagulation cascades, microglia cell activation, retina homeostasis, PUFA synthesis and sphingolipid metabolism. Blue representing low expression and red indicating higher levels. Also see supplementary figure S2.

RNAseq analyses of RPE from 14-mo mice showed significant alterations in genes implicated in multiple homeostatic pathways in *Grn^−/−^* mice RPE (Figures 4C-4E and Figure S2). KEGG pathway analysis identified complement C3 and complement and coagulation cascades pathway among the top 20 pathways significantly altered in *Grn^−/−^* RPE (Figure S2a). GO GSEA analyses showed upregulation of pathways associated with positive regulation of cell migration, inflammatory response, and microglia activation, pointing to an activated microglia-dependent inflammatory response in *Grn*^−/−^ mice (Figure 4C).

Further analysis of gene correlation revealed a strong association between complement and coagulation cascades genes (*C3, C1QA, C1QB, SERPINE1, C3AR1, ITGB2*) and microglia cell activation genes (*ITGAM, TREM2, TLR1*) (Figure S2B) and identified *C3, C3AR1, ITGAM, ITGAX,* and *ITGB2* as critical factors in regulating microglia activation (Figure 4D). The consequences of complement and microglia activation are reflected in the deregulation of pathways related to retinal homeostasis, sphingolipid metabolism, and polyunsaturated fatty acid synthesis in *Grn*^−/−^ mice (Figure 4E). Overall, these transcriptomic data suggest that mitochondrial and metabolic functions are compromised early in *Grn*^−/−^ RPE, which in turn drive complement signaling and microglial activation.

### High-speed super-resolution live imaging of RPE explants reveals early and widespread mitochondrial deficits in *Grn*^−/−^ mice RPE

Given the early transcriptional changes in mitochondrial ROS metabolism, we decided to investigate mitochondrial dynamics and function by high-speed super-resolution live imaging of RPE explants from wildtype and *Grn*^−/−^ mice ^30^. Live imaging of MitoTracker-labeled mitochondria and volumetric reconstructions of mitochondrial networks showed increased connectivity or hyperfusion of mitochondria in 4-mo *Grn*^−/−^ RPE that was even more pronounced in 17-mo mice (Figures 5A-5C).

**Figure 5.**
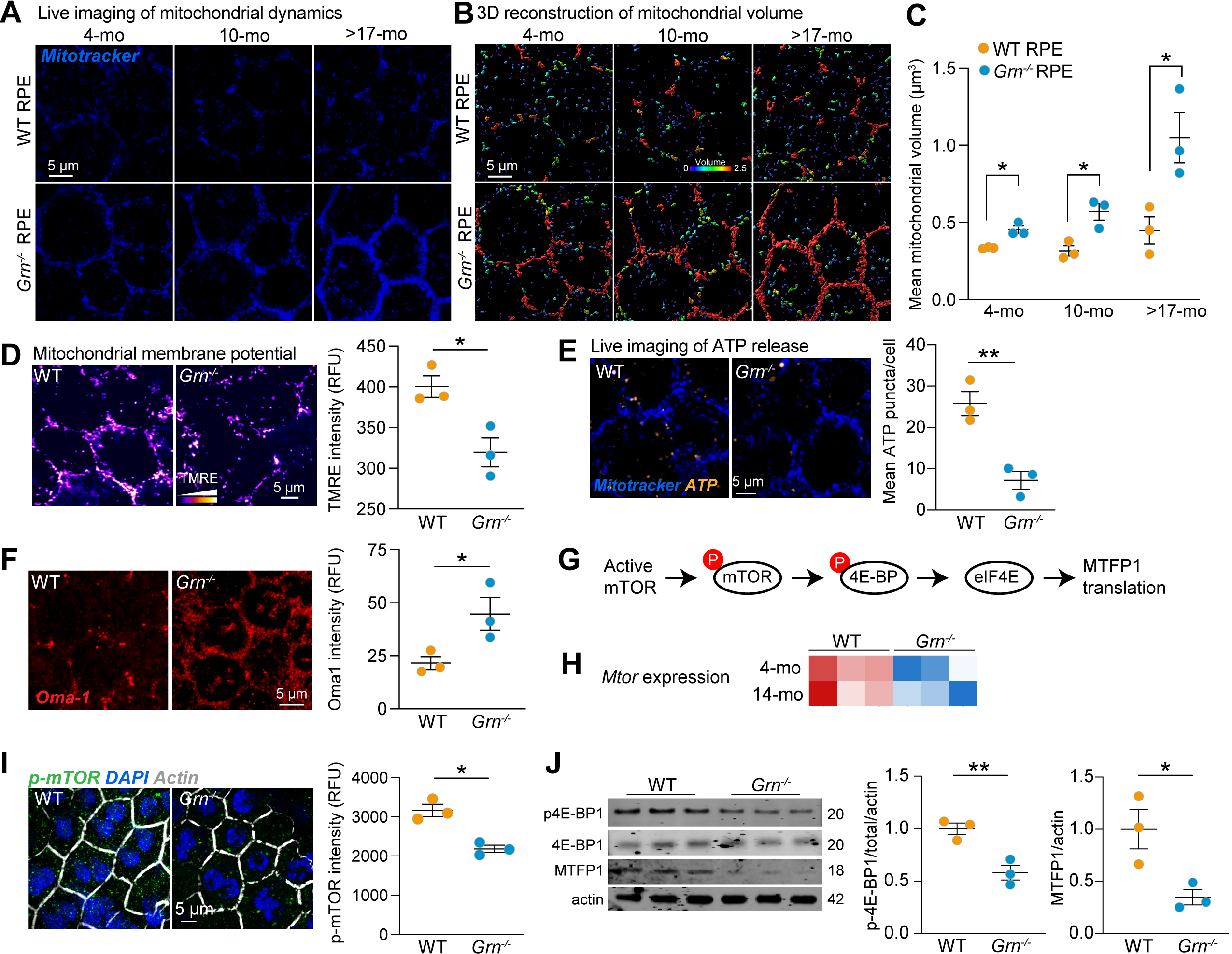
Widespread mitochondrial dysfunction in 4-month-old *Grn^−/−^* RPE mediated by inhibition of the mTOR-MTFP1 axis. (**A**) Stills from live imaging of mitochondrial dynamics in 4-, 10-, and >17-mo WT and *Grn^−/−^* mice RPE flatmounts using MitoTracker Deep Red. (**B**) Volumetric reconstuctions of mitochondrial volumes from live imaging in (A). Warmer colors: fused/integrated mitochondria; cooler colors: discrete mitochondria. (**C**) Quantification of mitochondrial volumes from live imaging data showing increased volumes (hyperfused mitochondria) in *Grn^−/−^* RPE. Mean ± SEM, 3 mice/genotype and age. *, *p* < 0.05 by two-way ANOVA with Tukey’s multiple comparisons test. (**D**) Live imaging stills and quantification of mitochondrial membrane potential (TMRE) in 4-mo WT and *Grn^−/−^* mice RPE flatmounts. Mean ± SEM, n =3. *, *p* < 0.05. (**E**) Live imaging stills and quantification of ATP release (BioTracker ATP Red) in 4-mo WT and *Grn^−/−^* mice RPE flatmounts. Mean ± SEM, n = 3. **, *p* < 0.01. (**F**) Oma1 immunostaining (red) and quantification in 4-mo WT and *Grn^−/−^* mice RPE flatmounts. Mean ± SEM, n =3. *, *p* < 0.05. (**G**) Schematic of mTOR/4E-BP1 signaling that regulates translation of MTFP1 (mitochondrial fission process 1). (**H**) Heatmaps showing decreased *Mtor* expression in 4-month-old *Grn^−/−^* RPE. (**I**) phospho-mTOR immunostaining (green) and quantification in WT and *Grn^−/−^* mice RPE flatmounts. Actin: gray and DAPI: blue. Mean ± SEM, n = 3. *, *p* < 0.05. (**J**) Immunoblotting of WT and *Grn^−/−^* mice RPE lysates with antibodies to phospho-4E-BP1, 4E-BP1, MTFP1, and β-actin and quantification. Mean ± SEM, n = 3. *, *p* < 0.05, **, *p* < 0.01. Also see supplementary figure S3.

We next evaluated the impact of this hyperfusion on mitochondrial function using tetramethyl rhodamine, methyl ester (TMRE) to measure mitochondrial membrane potential (Δψ_m_) and BioTracker ATP Red to measure mitochondrial ATP in live RPE explants. These studies revealed a pronounced loss of both Δψ and ATP generation (Figures 5D and 5E) indicative of bioenergetic deficits in *Grn*^−/−^ RPE.

Mitochondrial inner and outer membrane fusion events are regulated by the long forms of optic atrophy 1 (L-OPA1) and mitofusin 1 and 2 (MFN1 and MFN2), respectively. Mitochondrial outer membrane fission is orchestrated by the dynamin-related protein 1 (DRP1), which is activated by phosphorylation at Ser616 or dephosphorylation at Ser637, and the mitochondrial fission protein FIS1 ^31,32^. Dysregulation of these machinery, such as increased degradation of DRP1, has been shown to promote mitochondrial hyperfusion ^33^. However, immunoblotting showed no discernible differences in the expression of L-OPA1, MFN2, Ser616 and Ser637 DRP1, and FIS1 between wildtype and *Grn*^−/−^ RPE (Figure S3). In contrast, levels of the stress-activated mitochondrial protease Oma1 ^34,35^ were significantly elevated in *Grn*^−/−^ RPE (Figure 5F). Decreased Δψ_m_, increased ROS, or loss of ATP can stabilize Oma1, which cleaves a variety of substrates such as OPA1 (optic atrophy 1), PGAM5 (phosphoglycerate mutase 5), PINK1 (PTEN-induced kinase 1), and DELE1 (DAP3 Binding Cell Death Enhancer 1) to modulate mitochondrial fission/fusion, mitophagy, and the integrated stress response ^36^. Thus, multiple lines of evidence from transcriptomic, live imaging, and biochemical data strongly indicate a widespread impairment of mitochondrial structural and functional integrity in *Grn*^−/−^ RPE.

### A defective mTOR-MTFP1 signaling axis could drive mitochondrial hyperfusion in *Grn^−/−^* RPE

To uncover the mechanism of mitochondrial hyperfusion in *Grn*^−/−^ RPE, we focused on Mitochondrial Fission Process 1 (MTFP1, also known as MTP18), an inner mitochondrial membrane protein crucial for mitochondrial fission ^37,38^. Inhibition of the mechanistic target of rapamycin (mTOR) has been reported to induce mitochondrial hyperfusion by decreasing 4E-BP1-mediated MTFP1 translation in mouse embryonic fibroblasts (Figure 5G) ^39^. Transcriptomic and immunostaining data revealed downregulation of mTOR at the transcript and protein levels in 4- and 14-mo *Grn^−/−^*RPE compared to age-matched wildtypes (Figures 5H and 5I). Consequently, phosphorylated 4E-BP1 (p4E-BP1) and MTFP1 protein levels were significantly lower in *Grn^−/−^* RPE (Figure 5J), confirming that mTOR inhibition indeed decreases MTFP1 expression. Taken together, these studies identify a novel mechanism mediated by mTOR and MTFP1 that drives early mitochondrial hyperfusion and functional deficits in *Grn^−/−^*RPE.

### RPE-microglia crosstalk facilitates complement C3a-C3a receptor signaling in *Grn^−/−^* mice retinas

We next asked whether mitochondrial hyperfusion in *Grn^−/−^*RPE induces migration of microglia into the subretinal space by modulating complement activity, given the upregulation of complement pathway genes in 4-month-old *Grn^−/−^* RPE. Mitochondrial hyperfusion has been shown to activate NF-κ;B ^40^, to induce expression of the complement protein C3 and the C3a receptor (C3aR) ^41^. In the retina, a low level of complement activation in the retina controls the balance between inflammation and immune response. The RPE is a major biosynthetic source of complement proteins including C3 in the healthy retina ^42,43^. Under conditions of cell stress, we and others have reported that full-length C3 undergoes intracellular proteolysis by cathepsin L to release ∼8 kDa biologically active C3a peptides ^44–46^. C3a-C3aR signaling activates STAT3, which regulates the expression of target genes implicated in microglial activation, mitochondrial fusion, and inflammation ^47–49^.

RNAseq analyses confirmed upregulation of canonical NF-κ;B target genes and increased C3 and C3aR expression in 4- and 14-mo *Grn^−/−^* RPE compared to age-matched wildtypes (Figures 6A and 6B). Consistent with overactive C3a-C3aR signaling, several STAT3 target genes were also upregulated in *Grn^−/−^* RPE (Figure 6C). Quantification of C3 and C3a levels in 4-mo mice RPE flatmounts by immunostaining with specific antibodies ^45^ showed that processing of C3 to biologically active C3a was significantly elevated in *Grn^−/−^* RPE (Figure 6D), likely due to upregulation of cathepsin L (Figure 4B).

**Figure 6.**
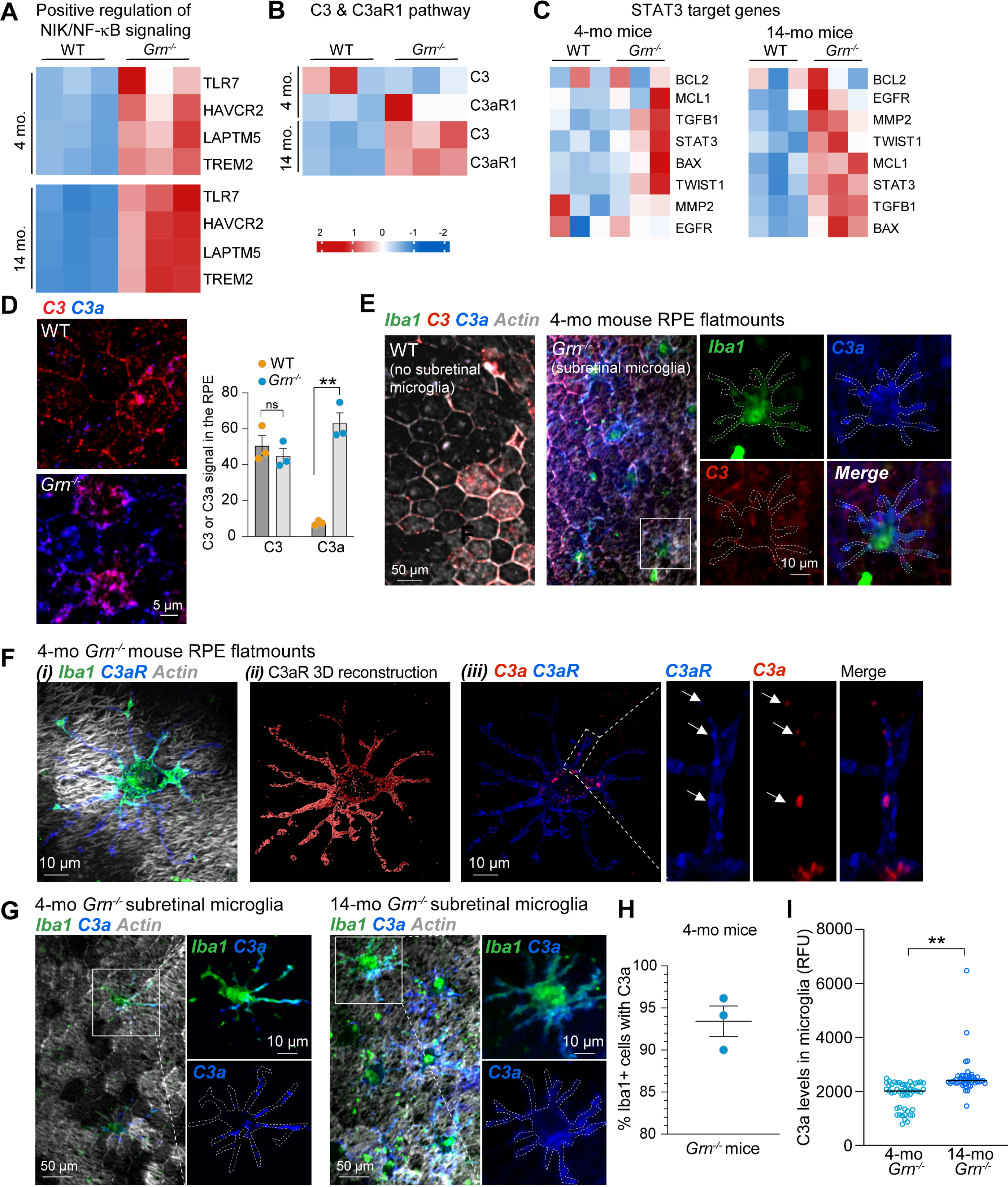
RPE-microglia crosstalk facilitates complement C3a-C3a receptor signaling in *Grn^−/−^* mice retinas. Heatmaps of (**A**) NIK/NF-kB regulated genes, (**B**) positive regulation of C3/C3aR pathway genes, and (**C**) STAT3 target genes in 4- and 14-mo WT and *Grn^−/−^* RPE. (**D**) Immunostaining and quantification of C3 (red) and C3a (blue) levels in 4-mo WT and *Grn^−/−^* RPE. Mean ± SEM, n =3. *, *p* < 0.01, two-way ANOVA with Tukey’s multiple comparisons test. ns – not significant. (**E**) Representative images of 4-mo WT and *Grn^−/−^* RPE flatmounts immunostained for Iba1 (green), C3 (red), and C3a (blue). Only *Grn^−/−^* RPE flatmounts have subretinal microglia, which accumulate high levels of C3a (high-magnification images of regions demarcated in white squares). (**F**) *(i)* Representative high-magnification image of a sub-retinal microglia on 4-mo *Grn^−/−^*RPE flatmount immunostained for Iba1 (green), C3a (red), and C3aR (blue). *(ii)* Volume reconstruction of C3aR (red) on the microglial surface. *(iii)* Colocalization of C3a (red) and C3aR (blue) in microglial processes (white arrows). (**G**) Immunostaining of C3a (blue) in subretinal microglia (green) of 4- and 14-mo *Grn^−/−^* RPE. Quantification of (**H**) % Iba1-positive subretinal microglia that had C3a in 4-mo *Grn^−/−^* mice and (**I**) C3a immunostaining intensity in Iba1+ subretinal microglia in 4- and 14-month-old *Grn^−/−^*mice. Mean ± SEM, n =3. ** *p* < 0.01.

In the brain, microglia are the major cells that express C3aR and aberrant C3-C3aR activation has been implicated in promoting neurodegeneration in mouse models of Alzheimer’s disease ^47,50^. In *Grn^−/−^* retinas, immunostaining for C3, C3a, and C3aR revealed that the majority (∼93%) of Iba1+ subretinal microglia had abundant C3a but very low C3 (Figures 6E, 6G, and 6H), suggesting that microglia likely take up C3a released by the underlying RPE, thereby activating C3aR. High-resolution immunostaining and 3D volume reconstructions showed that C3aR was enriched on the cell surface of subretinal microglia and partially colocalized with C3a (Figure 6F). Microglia in 14-mo *Grn*^−/−^ mice accumulated significantly more C3a compared to 4-mo *Grn*^−/−^ mice (Figures 6G and 6I), indicating that progressive C3-C3aR-dependent microglial activation could contribute to retinal degeneration in this model.

### C3aR antagonism corrects RPE mitochondrial deficits and limits the pro-inflammatory phenotype of subretinal microglia in young *Grn^−/−^* mice

To establish whether blocking C3aR signaling could ameliorate some of the early retinal pathology mediated STAT3 activation, we administered SB290157, a C3aR antagonist (C3aRA, 0.5 mg/kg ^51,52^) or vehicle (PBS) intraperitoneally to 1-month-old *Grn*^−/−^ mice for 8 weeks. STAT3 activates transcription of genes involved in mitochondrial fusion ^48,53^ and has been implicated in driving microglia to a proinflammatory phenotype ^54,55^. As STAT3 is a direct target of C3aR ^47^, we reasoned that inhibiting C3aR could potentially target both mitochondrial hyperfusion and activation of subretinal microglia in *Grn*^−/−^ mice. In support of this, immunostaining and quantification of mitochondrial volumes in RPE flatmounts showed that C3aRA treatment normalized mitochondrial networks in *Grn*^−/−^ mice RPE (Figures 7A and 7B). The number of subretinal microglia with C3a (Figures 7C and 7D) and CD68 (Figures 7E and 7F) were significantly decreased after C3aRA administration compared to vehicle-treated *Grn*^−/−^ mice, suggesting that C3aR antagonism could limit microglia-mediated neuroinflammation. Collectively, these data support a mechanistic link between RPE mitochondrial hyperfusion and subretinal microglial activation via aberrant intercellular C3a-C3aR signaling in models of progranulin deficiency.

**Figure 7.**
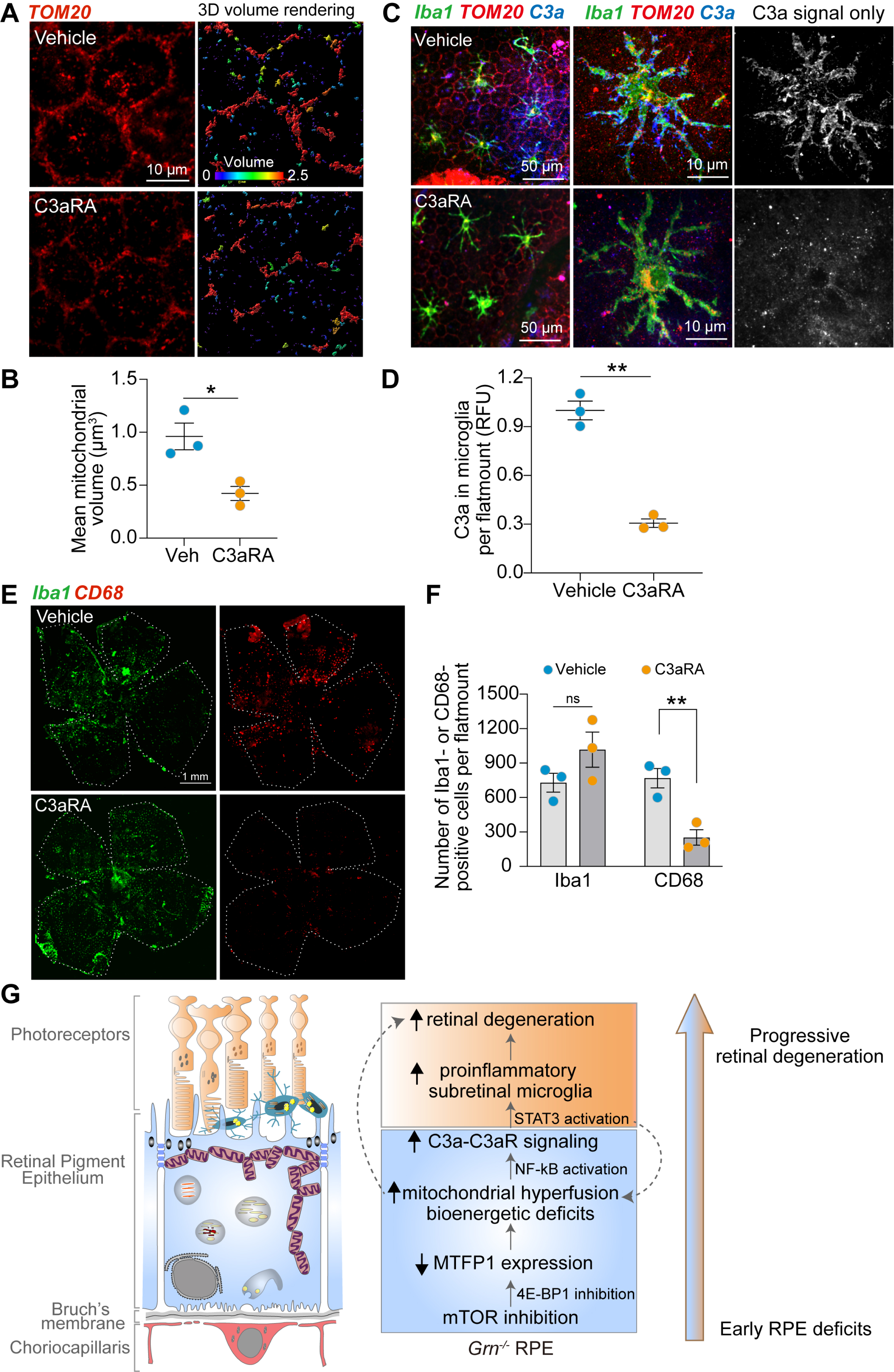
C3aR antagonism corrects RPE mitochondrial deficits and limits the pro-inflammatory phenotype of subretinal microglia in young *Grn^−/−^* mice. (**A**) TOM20 (red) immunostaining and mitochondrial volume reconstructions in RPE flatmounts from vehicle- or C3aRA-treated 3-mo *Grn^−/−^* mice. (**B**) Quantification of RPE mitochondrial volumes. Mean ± SEM, n = 3 mice per treatment. * *p* < 0.05. (**C**) Low- and high-magnification images of subretinal microglia immunostained for Iba1 (green), C3a (blue), and TOM20 (red). Single channel images of zoomed in microglia show C3a in gray. (**D**) Quantification of microglial C3a levels per RPE flatmount in vehicle- or C3aRA-treated *Grn^−/−^* mice. Mean ± SEM, n = 3 mice per treatment. ** *p* < 0.01. (**E**) Representative RPE flatmounts from vehicle- or C3aRA-treated *Grn^−/−^* mice. Flatmounts were immunostained for Iba1 (green) or CD68 (red). (**F**) Quantification of numbers of Iba1- or CD68-positive subretinal microglia in vehicle- or C3aRA-treated *Grn^−/−^* mice. Mean ± SEM, n = 3 mice per treatment. ** *p* < 0.01. ns – not significant. (**G**) Schematic representation of mechanisms by which progranulin loss drives mitochondrial dysfunction in the RPE, C3a-C3aR signaling, and subretinal microglial activation.

## DISCUSSION

The retina is a uniquely accessible extension of the CNS as it shares developmental and anatomical features with the brain and is directly connected to it via the optic nerve ^56^. This, coupled with the development of advanced non-invasive imaging technologies that can monitor retinal structure and function, makes the retina a powerful platform to interrogate neurodegenerative diseases. Indeed, retinal imaging and histopathological studies have identified changes in the retinal ganglion cell layer, retinal vasculature, and the optic nerve in patients with mild cognitive impairment and early-stage Alzheimer’s disease ^57–61^. In this study, using semi-automated OCT segmentation, we observed thinning of the photoreceptor outer nuclear layer and thickening of the outer plexiform layer in *GRN* mutation carriers compared to healthy controls, likely indicative of photoreceptor loss and microglial expansion. Longitudinal assessments showed that *GRN* mutation carriers with higher OPL/ONL ratios exhibited greater cognitive impairment and increased white matter abnormalities over time. ONL thinning and OPL expansion have also been reported in patients with FTD-tauopathy (FTD-Tau) ^62,63^, suggesting that these changes could be powerful biomarkers to diagnose FTD and monitor disease progression.

Progranulin has been implicated in modulating inflammation and recent studies suggest that in the brain, progranulin loss transitions microglia from a homeostatic state to a disease-activated, hyperinflammatory state ^64^. This shift leads to severe microglial pathology, including impaired phagocytosis ^65^ and accumulation of myelin debris in microglial lysosomes ^66^. Activation of microglia in the thalamus of *Grn*^−/−^ mice brains has been reported after 7 months with neuronal loss occurring over a span of 18-23 months ^14,67^. Compared to the brain, retinal neurodegeneration occurred on an accelerated timespan in *Grn*^−/−^ mice: we observed infiltration of microglia from the IPL/OPL into the subretinal space as early as 3 months and significant photoreceptor degeneration at 10 months, suggesting that *Grn*^−/−^ mice recapitulate early retinal pathology observed in *GRN* mutation carriers.

To dissect how progranulin deficiency promotes retinal degeneration and identify the cellular and molecular players involved, we conducted multimodal analyses of wildtype and *Grn*^−/−^ mice retinas using transcriptomics, advanced live imaging, and biochemical approaches (Figure 7G). As both full-length progranulin and granulin peptides generated by its proteolysis have been implicated in cathepsin D expression and activation ^7,8^, we initially hypothesized that loss of cathepsin D function in *Grn*^−/−^ RPE would inhibit degradation of phagocytosed outer segment proteins, which could explain lipofuscin accumulation in the retinas of *Grn*^−/−^ mice and *GRN*-FTD patients ^13,20^. However, both cathepsin D expression and phagolysosomal degradation of outer segments are unimpaired in *Grn*^−/−^ RPE. This is not surprising because efficient outer segment phagocytosis by the RPE is fundamental to photoreceptor health and vision and the RPE has robust mechanisms to safeguard this process even under conditions of significant cellular stress ^68^. Providing further proof of this, high-resolution imaging revealed that subretinal microglia had phagocytosed photoreceptor outer segments and that it was these microglia, and not the RPE, that were the source of age-dependent lipofuscin autofluorescence.

Transcriptomic analyses provided important clues regarding potential mechanisms responsible for retinal injury in *Grn*^−/−^ mice: first, in contrast to global changes in lysosomal pathway genes observed in the brain ^22,69^, only a subset of lysosomal hydrolases was altered in *Grn*^−/−^ RPE, suggesting that lysosomal dysfunction is likely not the primary driver of retinal degeneration.

Second, our data showed a specific upregulation of genes involved in ROS metabolic processes and the complement pathway, indicating that mitochondrial injury and inflammation might be key pathological mechanisms in *Grn*^−/−^ RPE. To obtain a real-time assessment of mitochondrial form and function, we performed super-resolution live imaging of RPE explants because intricate mitochondrial networks are poorly preserved in fixed tissue ^30^. Unexpectedly, 4D analysis of live imaging data showed that mitochondria in 4-month-old *Grn*^−/−^ mice RPE were hyperfused with dissipated membrane potential and defective ATP production, indicative of a global disruption of mitochondrial homeostasis and bioenergetic deficits. Recent studies have reported a role for progranulin in mitochondrial biogenesis and mitophagy in podocytes, neuroblastoma cell lines, regulatory T-cells, and vascular smooth muscle cells, but insight into underlying mechanisms has been lacking ^70–73^.

How might progranulin loss compromise mitochondrial structural and functional integrity? Our studies show that mitochondrial hyperfusion is a consequence of decreased MTPF1 translation, a protein regulated by mTOR ^39^. Consistent with previous reports that progranulin activates the Akt-mTOR pathway ^74,75^, our transcriptomic analysis revealed a nearly 50% decrease in mTOR mRNA levels in 4-month-old *Grn*^−/−^ RPE, suggesting that mitochondrial hyperfusion induced by progranulin deficiency occurs via the mTOR pathway and is initiated early and rapidly. These data also identify MTFP1 as a critical regulator of mitochondrial dynamics in the RPE, because none of the other proteins that are known to modulate mitochondrial fission or fusion were altered in *Grn*^−/−^ RPE. Mitochondrial hyperfusion activated transcription of NF-kB target genes including complement protein C3 and the C3a receptor ^40,41^ in 4-month-old *Grn*^−/−^ RPE. C3 is intracellularly proteolyzed to C3a by cathepsin L ^46^, which was transcriptionally upregulated in *Grn*^−/−^ mice.

Patients with *GRN*-FTD have been reported to have increased levels of serum complement proteins ^76^. Upregulation of complement pathway genes (*C1QA, C1QB, C1QC,* and *C3*) and microglia activation genes (*CD68* and *TREM2*) has been reported in brain microglia of 16-month-old *Grn*^−/−^ mice. Consistent with this, we observed an early and sustained increase in complement and microglia genes in *Grn*^−/−^ mice retinas. Analysis of gene correlation and STRING interaction networks identified *C3, C3AR1*, and *ITGAM* as key players in microglial activation in the retina. The central role for C3a-C3aR signaling in modulating microglial homeostasis was supported by the striking increase in C3a in the RPE and C3aR on the surface of subretinal microglia in 4-month-old *Grn*^−/−^ mice.

STAT3, a downstream target of C3aR signaling, is a transcriptional activator of mitochondrial fusion machinery and has been implicated in reprogramming microglia from a homeostatic, neuroprotective phenotype towards a proinflammatory, neurotoxic state in experimental models of Parkinson’s disease, Alzheimer’s disease, diabetic retinopathy, and ocular hypertension ^54,55,77–80^. Our studies confirmed the role of the C3a-C3aR-STAT3 axis in mediating retinal pathology as pharmacological antagonism of C3aR in young *Grn*^−/−^ mice restored both RPE mitochondrial structural integrity and microglial homeostasis. Independent of its effects on microglial phenotypes, that C3aR antagonism corrects RPE mitochondrial deficits in *Grn*^−/−^ mice makes it a target worthy of further investigation in FTD and other neurodegenerative diseases. The RPE relies solely on oxidative phosphorylation for its energy requirements and spares glucose from the choroidal circulation for photoreceptors. Deficits in OXPHOS force the RPE to become more glycolytic, which starves photoreceptors, leading to retinal degeneration ^19,81,82^. C3aR antagonism could therefore exert a synergistic beneficial effect on the two main drivers of retinal degeneration caused by progranulin loss: RPE mitochondria and subretinal microglia.

Progranulin deficiency has been implicated in the pathogenesis of not only FTD and CLN11, but other neurodegenerative diseases including Alzheimer’s disease, Parkinson’s disease, and Limbic-predominant age-related TDP-43 encephalopathy (LATE) ^4,83^. Several of these diseases exhibit retinal phenotypes, but insight into the underlying mechanisms has remained largely elusive. Our studies now identify a novel stepwise feed-forward mechanism that directly connects progranulin loss with mitochondrial hyperfusion in the RPE, which leads to C3a/C3aR-mediated microglial activation in the subretinal space and progressive retinal degeneration (Figure 7G). The early manifestation of these changes in the retina compared to those observed in the brain suggests that progranulin deficiency has an accelerated and pronounced impact in derailing RPE homeostasis and activating retinal microglia. Our study underscores the significance of retinal changes as early indicators of neurodegenerative processes associated with *GRN* deficiency and identifies critical mechanisms that can be targeted to mitigate retinal pathogenesis in *GRN*-related disorders.

## STAR METHODS

### Patient cohort

We selected research participants with genetic FTLD from cohorts enrolled in longitudinal research at the University of California, San Francisco Memory and Aging Center (MAC UCSF). Inclusion criteria were (i) presence of heterozygous *GRN* mutation, (ii) macular and/or peripapillary OCT data and (ii) no neuro-ophthalmological or ophthalmological comorbidities influencing OCT results. Healthy control subjects (HC) were selected from the Hillblom Healthy Aging Study at the UCSF MAC. Subjects with neuro-ophthalmological or ophthalmological comorbidities potentially influencing OCT results were excluded (such as glaucoma, diabetic retinopathy and severe central serous chorioretinopathy). Research participants at the UCSF MAC undergo a comprehensive multidisciplinary assessment that includes functional, neuropsychological, and socioemotional assessments, as previously described ^84^. Cognitive data were available for 23 heterozygous *GRN* mutation carriers and 85 HCs cross-sectionally as well as 12 heterozygous *GRN* mutation carriers longitudinally with a follow-up of [median(min-max)] 28 (12-73) months. All HCs were tested for and did not carry any of the three FTLD genes including *GRN*. We included all cognitive testing acquired at and after the time of the first OCT. The study was approved by the local ethics committees and conducted in accordance with the applicable laws and the current version of the Declaration of Helsinki. All participants gave written informed consent. Data are reported according to STROBE reporting guidelines.

### Optical coherence tomography

All OCT data were acquired with the same Spectralis SD-OCT device (Heidelberg Engineering, Heidelberg, Germany). Peripapillary retinal nerve fiber layer thickness (pRNFL) was measured with activated eye tracker using 12* ring scans around the optic nerve head (3≤ART≤100). All other layers were measured using a 3.45mm diameter cylinder around the fovea from a macular volume scan (13 ≤ ART ≤ 72, 15 ≤ quality ≤ 42). OCT data reading and quality control was performed by three experienced raters (AK, SCM, FCO); image quality was assessed by OSCAR-IB criteria and reported according to APOSTEL recommendations ^85,86^. We excluded OCT scans with insufficient image quality from the quantitative analysis. Semi-automatic intraretinal layer segmentation was performed twice by two experienced raters (for confirmation, FCO and AK) using the software provided by the OCT manufacturer (Eye Explorer 1.10.4 with viewing module 6.12; Heidelberg Engineering). Qualitative OCT data analyses were executed by or under supervision of a board-certified neuro-ophthalmologist (AJG).

### Magnetic resonance tomography

Magnetic Resonance Imaging (MRI) data were available for 18 heterozygous *GRN* mutation carriers cross-sectionally and nine heterozygous *GRN* mutation carriers longitudinally with a follow-up of [median(min-max)] 1.4 (0.9-4.0) years. Participants were scanned on either a 3 Tesla Siemens Prisma fit or TrioTim at the UCSF Neuroscience Imaging Center. White matter hyperintensities were quantified on FLAIR-imaging using the Fazekas scale with a Fazekas score of 3 representing severe white matter disease – the low sample size for a Fazekas score of 3 limits the generalizability of this exploratory analysis ^25^.

### Volumetric processing

Prior to pre-processing, all T1-weighted images were visually inspected for quality control. Images with excessive motion or image artifact were excluded. Tissue segmentation was performed using SPM12 (Wellcome Trust Center for Neuroimaging, London, UK, http://www.Fil.ion.ucl.ac.uk/spm) unified segmentation ^87^. Total intracranial volume (TIV) was calculated for each subject as the sum of the gray matter, white matter, and cerebrospinal fluid segmentation.

### Analysis of clinical data

Statistical analyses were performed with R 3.6.1. Figures were created using R and Adobe Illustrator. For all calculations, statistical significance was established at p < 0.05. Group differences were tested with x^2^ for sex and Wilcoxon-Mann-Whitney U test for age. Cross-sectional and longitudinal group differences in OCT and cognition were evaluated by t-test and linear mixed effect models accounting for within-subject inter-eye dependencies as a random effect (for OCT parameters) as well as for age and sex in non-matched subsets respectively. OPL/ONL-ratio (OPL divided by ONL) was not set as an outcome parameter a priori but included as a combined outcome parameter with potential future as prognostic biomarker.

### Mice

Wildtype and *Grn*^−/−^ mice on C57BL/6J background ^5,21,88,89^ were raised under 12-h cyclic light on a standard diet. In some studies, the C3aR inhibitor SB290157 was administered intraperitoneally (0.5 mg/kg) to 1-month-old mice 3 times/week for 8 weeks. Depending on the experimental paradigm, mice of various ages were euthanized using CO_2_ asphyxiation, eyes were removed, and eyecups were processed for immunohistochemistry, live imaging, immunoblotting, or transcriptomics as previously detailed ^30,45,68,90,91^. All studies were approved by University of California, San Francisco animal care and use authorities and in accordance to ARRIVE guidelines.

### Measurement of retinal layers in mice

After enucleation, eyes were immersed in a cold fixative solution (4% PFA, 2.5% glutaraldehyde, and 0.1 M cacodylate buffer) for ∼48 hours. Eyes were then dehydrated in graded ethanol and were embedded in Technovit 7100 glycol methacrylate. Embedded eyes were sectioned and stained with Hematoxylin and Eosin (H&E). Imaging was performed using an upright Axiophot microscope (ZEISS, Oberkochen, Germany) equipped with a 40X objective on a 12-MP Insight camera. To assess retinal morphology, the thickness of the outer nuclear layer (ONL) and outer plexiform layer (OPL) was quantitatively analyzed. Measurements were taken at a minimum of ten locations from the optic nerve head in four retinal sections obtained from four mice in each experimental group using the Spot software.

### Immunostaining of mice RPE flatmounts

Enucleated eyes were dissected to remove the anterior portion of the eyes including the lens. Four relaxing cuts were made on the eye cups prior to fixation in 4% PFA for 1 h at room temperature. The retinas were then clipped and removed. Each RPE flatmount was then blocked for 1 h at room temperature in 300 μL of 2% BSA in PBS supplemented with 0.1% Triton-X. After blocking, flatmounts were incubated in primary antibodies (see Table S1) diluted in 1% BSA in PBS at 4°C for 16h, followed by incubation in AlexaFluor secondary antibodies (1:500, ThermoFisher Scientific, Waltham, MA) and phalloidin (1:200, Cytoskeleton, PDHG1) diluted in 1% BSA for 2 h at room temperature. After secondary staining, flatmounts were rinsed as above and stained with DAPI (Sigma-Aldrich, St. Louis, MO; D9542, 1:200 in PBS for 15 min at room temperature). After three rinses, RPE flatmounts were mounted and sealed on clean slides with PBS and Vectashield (Vector Labs, Peterborough, UK). Images were acquired on Nikon spinning disk confocal microscope with 100x (1.49 NA) objective.

### Transcriptomics and bioinformatics analysis

Total RNA was isolated using the RNAqueous^TM^-Micro Total RNA Isolation Kit (Thermo Fisher, AM1931) immediately upon euthanasia. Anterior portion, the lens, and retinas were removed, and the remaining eye cups were kept on ice at all times. RPE were dissociated from the eyecups in sterile 1xPBS and lysis buffer provided in the kit was added to the cell suspension. Cell lysates were vortexed at maximum speed for 20s. Half-volume of molecular-grade ethanol was added followed by brief vortexing. Lysates were then passed through the Micro filter cartridge via 14,000x*g* centrifugation, followed by three washes using the wash solutions provided. RNA was eluted in the elution buffer provided and subjected to DNase I treatment. Samples were then submitted to Novogene (Sacramento, CA) for bulk RNA sequencing using Illumina NovaSeq6000 platform. All samples were confirmed to have RIN > 7 using the RNA Nano 6000 Kit of the Bioanalyzer 2100 system (Agilent Technologies, CA). FASTQ were processed through fastp, and paired-end clean reads were aligned to the reference genome using the Spliced Transcripts Alignment to a Reference (STAR) software (52). The p-values were adjusted using the Benjamini & Hochberg (BH) procedure, which was performed using the R Studio software. Gene ontology analysis was conducted on the selected genes using DAVID Bioinformatics Resources ^92,93^, and the enrichment was determined based on the p-value and gene ratio. The data were visualized using the hiplot database (https://hiplot.cn/basic) ^94^.

### Immunoblotting

RPE cells were collected from mouse eyecups immediately after enucleation. Cells were lysed in 1X HNTG lysis buffer (50 mM HEPES, 150 mM NaCl, 10% glycerol, 1.5 mM MgCl2, 1% Triton X-100) supplemented with a protease inhibitor cocktail. Protein concentrations were measured by the DC Protein Assay (Bio-Rad). NuPAGE LDS sample buffer (4X) was added to the lysates, and samples were heated to 85°C for 10 minutes. Samples were resolved on a NuPAGE 4-12% Bis-Tris gel (Invitrogen) at 150V for 1 hour at room temperature. Gels were transferred onto nitrocellulose membranes using the iBlot transfer device (Invitrogen) for 7 minutes using the preset program 3. Membranes were blocked in Intercept (PBS) Protein-Free Blocking Buffer (LI-COR) for 1 h at room temperature with gentle shaking and incubated with indicated primary antibodies diluted in blocking buffer supplemented with 0.1% Tween20. Membranes were then incubated with IRDye secondary antibodies (1:5000, LI-COR) for 1 h at room temperature and scanned using LI-COR Odyssey CLx. Band intensities were quantified using Image Studio (LI-COR Odyssey).

### Live imaging of mitochondrial integrity and function

Freshly harvested mouse RPE flatmounts were dissected into leaflets and placed in DMEM supplemented with 1% FBS and 1% NEAA. Leaflets were labeled with Mitotracker Deep Red (300 nM, 15 min), BioTracker ATP Red (1:1000, 15 min), TMRE (500 nM, 10 min), and/or Cell Mask Green (1.5 μL/mL, 15 min), diluted in Fluorobrite DMEM, at 37℃. After staining, leaflets were rinsed, mounted onto a Warner chamber in Fluorobrite DMEM, and imaged immediately. Imaging was performed at 37°C in an Okolab humidified microenvironmental chamber on the Nikon spinning disk confocal microscope using a CFI60 Apochromat TIRF 100X oil immersion objective (1.49 NA), the Nikon Live-SR super-resolution module, and an Andor iXon Ultra 888 EMCCD camera. Identical laser power, exposure and gain settings were applied for all conditions within a set of experiments. in an Okolab humidified chamber at 37℃ on Nikon spinning disk confocal microscope using 100x (1.49NA) objective. Imaging duration for each leaflet was kept to less than 30 min to avoid artefacts. Acquired images were analyzed using the “Surfaces” module in Imaris (Bitplane) as previously detailed ^30,68^.

### Image analysis

All images were subjected to Gaussian filtering and background subtraction in Imaris (Bitplane). 3D-reconstructions of MitoTracker, TMRE, and BioTracker ATP signals were performed using the “Surfaces” tool with the same threshold applied to all images within an experiment ^30,68^. Mitochondrial volumes, mean TMRE intensities, and number of ATP surfaces (normalized to cell number) were obtained from the statistics tab and compiled in Microsoft Excel. For outer segment phagosome quantification, 3D reconstructions were performed as above for the rhodopsin channel, and the number of surfaces were recorded as the number of phagosomes. For quantifying microglia, the “Spots” tool was used and manual correction was performed as necessary. For analysis of microglia autofluorescence and TMEM119 intensities, 3D reconstructions were performed using the Iba1 channel and the mean intensities of the indicated channels within these IBA1 surfaces were recorded and compiled in Microsoft Excel. Background intensities were defined as mean of intensities from regions devoid of Iba1+ cells, using the phalloidin channel for surface creation. Graphs and statistical analyses were done using GraphPad Prism.

### Statistics

Statistical analyses of in vivo data were performed using Prism 9 (GraphPad). Details of all statistical tests, sample sizes, and statistical significance are reported in the figure legends. Unless otherwise noted in the legend, data are presented as means and error bars represent SEM. Statistical significance was defined as *p*-value < 0.05.

## Supplementary materials

Data Tables S1-S6

Supplementary Figures S1-S3

STAR Methods

Table S7. Key Resources

References

## Authors contribution

L.X.T., F.C.O., A.C., F.M.E., and A.L. developed the concept, planned, and performed experiments, analyzed data. L.X.T. and A.C. performed in vitro and in vivo biochemical analyses, immunofluorescence, imaging assay and experiments on *Grn*^−/−^ mice. L.X.T. performed RNA-seq and live imaging experiments. A.C. carried out immunoblot analysis, bioinformatics analysis and C3aR antagonist studies. F.C.O., K.Y., Y.C., A.K., C.F., D.J.B., A.A., A.S., S.C.M., M.C., R.Y.C., C.C., M.E.W., K.C., J.H.K., H.J.R. and B.L.M. collected and analyzed clinical data for the study. L.X.T., F.C.O., and A.C. performed biostatistical analyses. F.C.O, L.X.T, A.C, and A.L. made the figures. A.C. and A.L. wrote the manuscript with input from L.X.T and F.C.O. L.X.T., F.C.O., A.C., F.M.E., and A.L. edited the manuscript. F.M.E. and A.L. supervised the project, provided the resources, and acquired funding. All authors approved the final version of the manuscript.

## Data and materials availability

All data associated with this study are presented in the article and or the Supplementary Materials. RNAsequencing data will be uploaded to the NCBI Gene Expression Omnibus (GEO) repository (GSe). Only de-identified patient data is presented in the manuscript. All mouse lines and reagents used in this study are available commercially and will be shared upon request.

