## Supplemental information for "Targeting complement C3a receptor resolves mitochondrial hyperfusion and subretinal microglial activation in progranulin-deficient frontotemporal dementia"

**This PDF file includes:**

Data Tables S1 to S5

Supplementary Figures S1 to S3 with legends

STAR Methods

Table S6. Key Resources

Supplementary References

### Tables

##### Table S1. Cohort description

| | Healthy controls | Heterozygous *GRN* mutation carriers |
|---|---|---|
| **Subjects [N]** | 85 | 23 |
| **Age [years; mean ± SD]** | 69 ± 15 | 62 ± 11 |
| **Gender [male/female]** | 42/43 | 11/12 |
| **CDR score [median (IQR)]** | 0 (0 – 0) | 0.5 (0 – 0.5) |
| **Education [years, mean ± SD]** | 17 ± 2 | 17 ± 4 |

Abbreviations: CDR: clinical dementia rating, GRN: progranulin, IQR: inter-quartile range, SD: standard deviation.

##### Table S2. Cross-sectional OCT differences between healthy controls and heterozygous *GRN* mutation carriers.

| | HC [mean ± SD] | Heterozygous *GRN* mutation carriers [mean ± SD] | Beta | S | p | $R^2$m | $R^2$c |
|---|---|---|---|---|---|---|---|
| **pRNFL** [µm] | 91.22 ± 12.44 | 89.42 ± 10.45 | - 3.167 | 2.669 | 0.238 | 0.146 | 0.921 |
| **TMV** [mm$^3$] | 3.09 ± 0.14 | 3.09 ± 0.13 | - 0.035 | 0.033 | 0.287 | 0.186 | 0.967 |
| **mRNFL** [mm$^3$] | 0.22 ± 0.02 | 0.22 ± 0.02 | - 0.001 | 0.006 | 0.821 | 0.015 | 0.800 |
| **GCIPL** [mm$^3$] | 0.78 ± 0.08 | 0.77 ± 0.07 | - 0.023 | 0.017 | 0.193 | 0.249 | 0.962 |
| **INL** [mm$^3$] | 0.33 ± 0.03 | 0.33 ± 0.03 | 0.001 | 0.007 | 0.889 | 0.053 | 0.849 |
| **OPL** [mm$^3$] | 0.30 ± 0.03 | 0.33 ± 0.05 | 0.031 | 0.009 | **< 0.001** | 0.108 | 0.628 |
| **ONL** [mm$^3$] | 0.69 ± 0.07 | 0.66 ± 0.07 | - 0.039 | 0.016 | **0.019** | 0.125 | 0.899 |
| **RPE** [mm$^3$] | 0.16 ± 0.01 | 0.15 ± 0.02 | - 0.006 | 0.003 | **0.039** | 0.062 | 0.716 |

*Abbreviations*: pRNFL: GCIPL: combined ganglion cell and inner plexiform layer volume, INL: inner nuclear layer volume, mRNFL: macular retinal nerve fiber layer volume, peripapillary retinal nerve fiber layer thickness, ONL: outer nuclear layer volume, OPL: outer plexiform layer volume, RPE: retinal pigment epithelium volume, TMV: total macular volume.

**Table S3. Cross-sectional differences in cognitive domains (z-scores) between heterozygous *GRN* mutation carriers with high versus low OPL/ONL-ratio.**

| | **Baseline values [mean ± SD] for** | | **t** | **95%-CI** | **p** |
|---|---|---|---|---|---|
| | **High OPL/ONL-ratio** | **Low OPL/ONL-ratio** | | | |
| **Executive** | 0.2 ± 0.5 | -0.1 ± 0.4 | 0.83 | -0.40; 0.81 | 0.436 |
| **Language** | 0.3 ± 0.5 | -0.4 ± 1.3 | 1.15 | -0.88; 2.24 | 0.307 |
| **Visuospatial** | < -0.1 | 0.2 ± 1.0 | -0.44 | -1.72; 1.25 | 0.682 |
| **Memory** | 0.1 ± 0.6 | -0.2 ± 1.5 | 0.43 | -1.55; 2.14 | 0.689 |

**Table S4. Longitudinal differences in cognitive function between heterozygous *GRN* mutation carriers with high versus low OPL/ONL-ratio. Reported statistics for group effect over time (interaction effect: group*follow-up).**

| | **Annualized change [mean ± SD)** | | **Beta** | **SE** | **p** | **$R^2$m** | **$R^2$c** |
|---|---|---|---|---|---|---|---|
| | **High OPL/ONL-ratio** | **Low OPL/ONL-ratio** | | | | | |
| **Executive** | | | | | | | |
| Composite z-score | -0.01 ± 0.37 | 0.29 ± 0.20 | 0.013 | 0.006 | **0.039** | 0.064 | 0.691 |
| Trails Time | 8.09 ± 15.35 | -4.50 ± 12.40 | -0.598 | 0.311 | 0.067 | 0.050 | 0.784 |
| Design Fluency | -0.09 ± 3.99 | 0.50 ± 2.89 | 0.076 | 0.062 | 0.226 | 0.021 | 0.625 |
| Digit span - backward | 0.18 ± 0.60 | 0.50 ± 1.29 | 0.023 | 0.014 | 0.115 | 0.047 | 0.830 |
| Verbal fluency - lexical | -2.60 ± 5.62 | 4.25 ± 3.50 | 0.273 | 0.080 | **0.002** | 0.122 | 0.744 |
| Stroop | -3.20 ± 17.16 | 12.00± 29.68 | 0.380 | 0.247 | 0.133 | 0.082 | 0.229 |
| Abstraction | -0.27 ± 0.65 | -0.67 ± 0.58 | 0.009 | 0.014 | 0.513 | 0.012 | 0.873 |
| **Language** | | | | | | | |
| Composite z-score | < 0.01 | < 0.01 | <0.001 | <0.001 | >0.99 | 0.295 | >0.99 |
| Boston naming test | 0.00 ± 0.00 | 0.50 ± 1.00 | 0.009 | 0.010 | 0.370 | 0.067 | 0.953 |
| Verbal fluency - semantic | -5.70 ± 9.76 | 0.00 ± 4.32 | 0.213 | 0.096 | **0.035** | 0.166 | 0.725 |
| **Face perception and socioemotional** | | | | | | | |
| CATS affect identification | 0.44 ± 1.24 | 2.00 ± 1.83 | 0.066 | 0.030 | **0.040** | 0.096 | 0.755 |
| **Visuospatial** | | | | | | | |
| Composite z-score | -0.19 ± 0.38 | -0.09 ± 0.50 | 0.003 | 0.006 | 0.577 | 0.054 | 0.845 |
| **Memory** | | | | | | | |
| Composite z-score | 0.08 ± 0.50 | -0.05 ± 0.20 | <0.001 | 0.008 | 0.960 | 0.028 | 0.839 |

**Table S5. Annualized atrophy in heterozygous *GRN* mutation carriers with high versus low OPL/ONL ratio**

| | Annualized atrophy rate [mean] for | | t | 95%-CI | p |
|---|---|---|---|---|---|
| | High OPL/ONL-ratio | Low OPL/ONL-ratio | | | |
| **Parietal lobe (left) [l]** | -0.001 | -0.001 | -0.544 | -0.003; 0.002 | 0.609 |
| **Parietal lobe (right) [l]** | -0.001 | 0.000 | -0.699 | -0.002; 0.001 | 0.511 |
| **Temporal lobe (left) [l]** | 0.000 | 0.000 | 0.151 | -0.001; 0.001 | 0.885 |
| **Temporal lobe (right) [l]** | 0.000 | 0.000 | 0.932 | -0.001; <0.001 | 0.392 |
| **Insula (left) [l]** | -0.003 | -0.013 | 0.826 | -0.023; 0.043 | 0.455 |
| **Insula (right) [l]** | -0.003 | -0.013 | 0.794 | -0.027; 0.048 | 0.476 |
| **Frontal lobe (left) [l]** | - 0.002 | -0.001 | -0.601 | -0.004, 0.003 | 0.570 |
| **Frontal lobe (right) [l]** | -0.001 | -0.001 | -0.701 | -0.003; 0.002 | 0.512 |
| **Occipital lobe (left) [l]** | 0.000 | -0.001 | 0.147 | -0.002; 0.002 | 0.888 |
| **Occipital lobe (right) [l]** | -0.001 | -0.001 | 0.206 | -0.002; 0.002 | 0.844 |

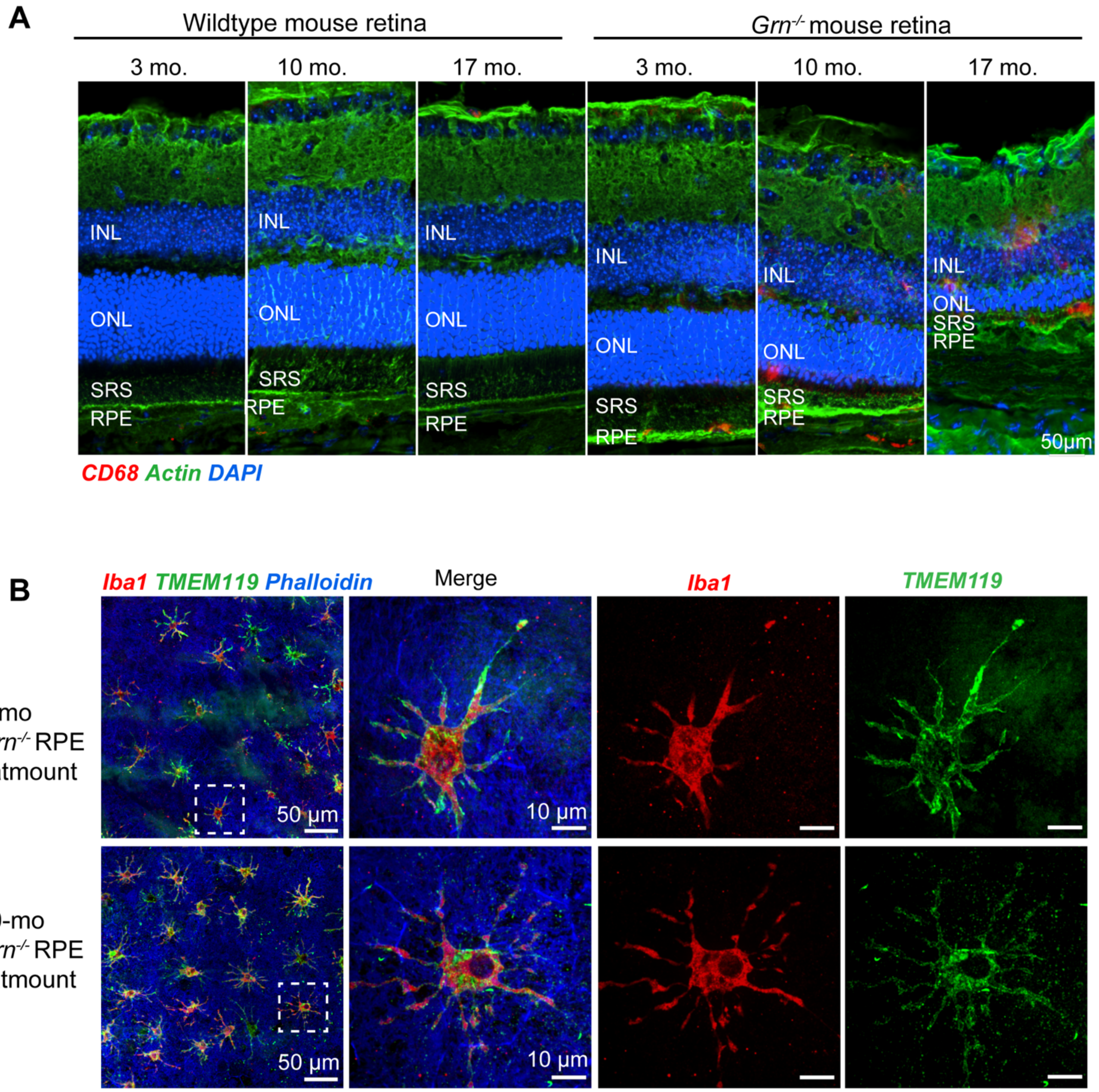

**Supplementary Figure S1. Progressive outer retinal degeneration in *Grn*$^{-/-}$ mice**. (**A**) Retinal cryosections from 3-, 10-, or 17-mo wildtype and *Grn*$^{-/-}$ mice immunostained for CD68 (red). Retinal layers are distinguished based on actin (green) and DAPI (blue) staining. RGC: Retinal ganglion cell layer; INL: Inner nuclear layer; OPL: Outer plexiform layer; ONL: Outer nuclear layer; SRS: Subretinal space; RPE: Retinal pigment epithelium. (**B**) TMEM119 (green) colocalization with Iba1 (red) in the subretinal space in 4- and 20-month-old *Grn*$^{-/-}$ mice.

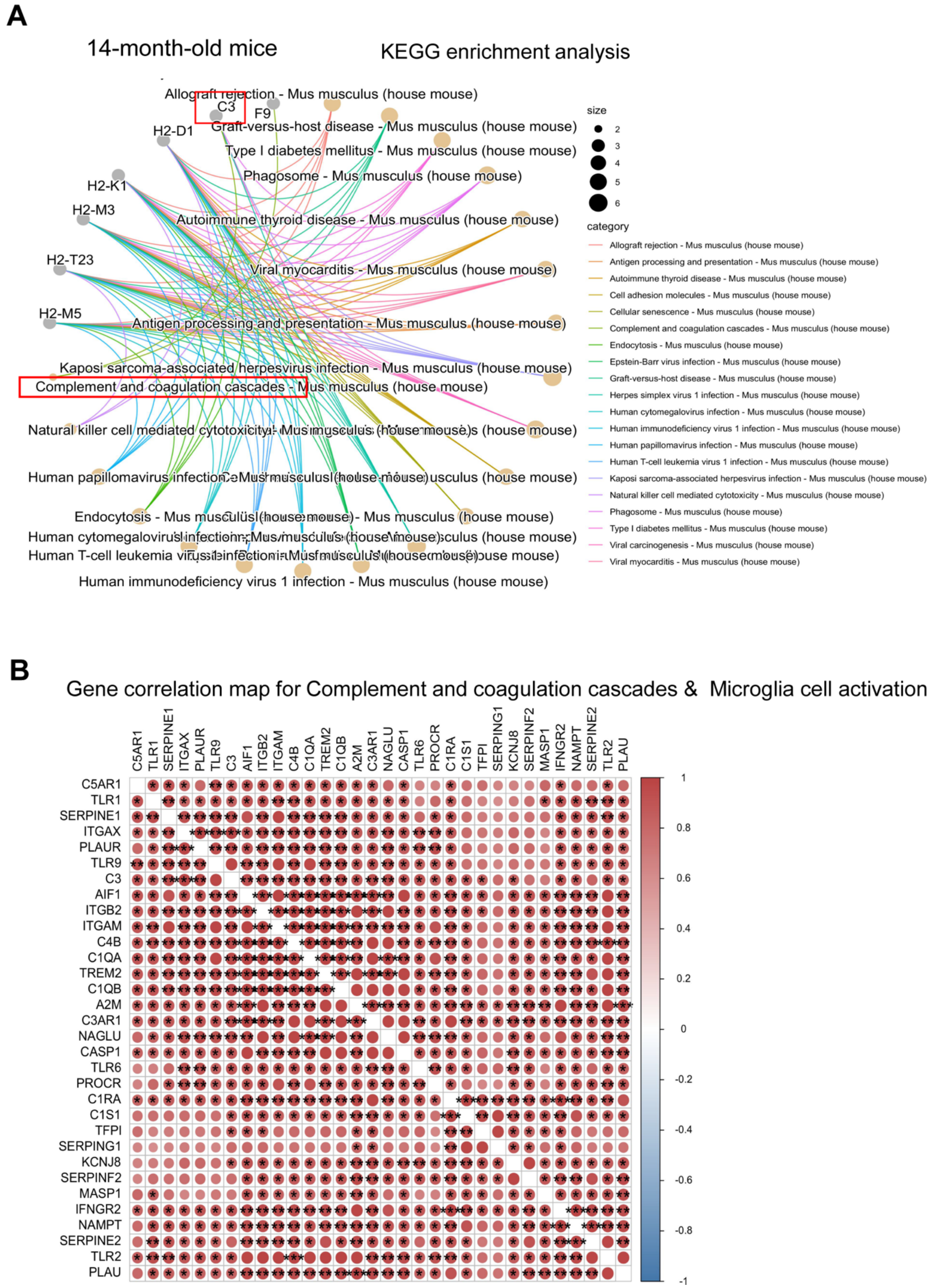

**Supplementary Figure 2. Activation of complement and microglial cascades in 14-month-old *Grn*$^{-/-}$ mice**.

(**A**) KEGG analysis of pathways enriched for genes that are differentially expressed in 14-month-old *Grn*$^{-/-}$ RPE relative to age-matched wildtypes, with the top 20 GO terms shown. (**B**) Gene correlation analysis for selected genes, revealing two specific interactions, marked with a star symbol.

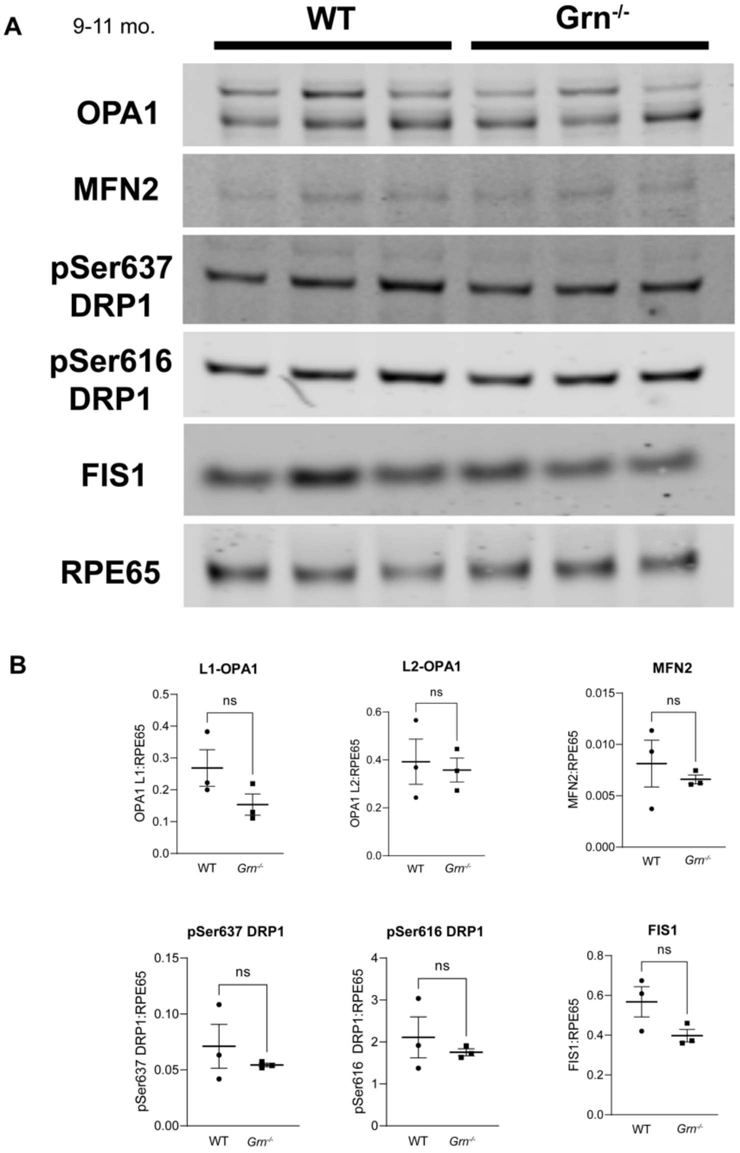

**Supplementary Figure 3. Expression of mitochondrial fusion and fission proteins in wildtype and *Grn*$^{-/-}$ RPE.** (A) Immunoblots and (B) quantification of indicated mitochondrial proteins in RPE lysates. RPE65 was used as the loading control. Mean ± SEM, n = 3 mice per genotype. n.s. – not significant, t-test.

#### EXPERIMENTAL METHODS

##### Patient cohort

We selected research participants with genetic FTLD from cohorts enrolled in longitudinal research at the University of California, San Francisco Memory and Aging Center (MAC UCSF). Inclusion criteria were (i) presence of heterozygous *GRN* mutation, (ii) macular and/or peripapillary OCT data and (ii) no neuro-ophthalmological or ophthalmological comorbidities influencing OCT results. Healthy control subjects (HC) were selected from the Hillblom Healthy Aging Study at the UCSF MAC. Subjects with neuro-ophthalmological or ophthalmological comorbidities potentially influencing OCT results were excluded (such as glaucoma, diabetic retinopathy and severe central serous chorioretinopathy). Research participants at the UCSF MAC undergo a comprehensive multidisciplinary assessment that includes functional, neuropsychological, and socioemotional assessments, as previously described [1]. Briefly, a functional assessment is done through a semi-structural interview with the patient's co-participant (caregiver) using the clinical dementia rating (CDR) [2]. Global cognition is assessed with the Mini-Mental State Examination (MMSE) [3]. Verbal and visual episodic memory are tested with the California Verbal Learning Test (CVLT) and a 10-minute delayed free recall of the Benson complex figure recall, respectively [4]. Visuospatial processing was evaluated with the Benson complex figure copy, and the Visual Object Space Perception (VOSP) [5]. Arithmetic calculations and praxis are also assessed [1]. Executive functioning was evaluated with backward digit span, Trails sequencing, Stroop color-naming test, and design fluency [6,7]. Word generativity was evaluated with lexical fluency (patients generated as many words starting with one letter in 60 seconds avoiding proper names) and semantic verbal fluency (patients generated as many animals as possible in 60 seconds) [6,7]. Verbal semantic knowledge was evaluated with the Peabody Picture Vocabulary Test (PPVT; patients were asked to choose the picture that best describes a word) [8], and the abbreviated 15-item Boston Naming Test (BNT; patients were asked to name different drawings) [9]. The ability to label emotional facial expressions with words is tested with the CATS emotion identification task, in which patients chose the emotion term that matched the facial expression depicted in a photograph from a list of multiple-choice options [10]. Cognitive data were available for 23 heterozygous *GRN* mutation carriers and 85 HCs cross-sectionally as well as 12 heterozygous *GRN* mutation carriers longitudinally with a follow-up of [median(min-max)] 28 (12-73) months. All HCs were tested for and did not carry any of the three FTLD genes including *GRN*. We included all cognitive testing acquired at and after the time of the first OCT. The study was approved by the local ethics committees and conducted in accordance with the applicable laws and the current version of the Declaration of Helsinki. All participants gave written informed consent. Data are reported according to STROBE reporting guidelines.

**Optical coherence tomography**

All OCT data were acquired with the same Spectralis SD-OCT device (Heidelberg Engineering, Heidelberg, Germany). Peripapillary retinal nerve fiber layer thickness (pRNFL) was measured with activated eye tracker using 12* ring scans around the optic nerve head ($3 \leq ART \leq 100$). All other layers were measured using a 3.45mm diameter cylinder around the fovea from a macular volume scan ($13 \leq ART \leq 72$, $15 \leq quality \leq 42$). OCT data reading and quality control was performed by three experienced raters (AK, SCM, FCO); image quality was assessed by OSCAR-IB criteria and reported according to APOSTEL recommendations [11]. We excluded OCT scans with insufficient image quality from the quantitative analysis. Semi-automatic intraretinal layer segmentation was performed twice by two experienced raters (for confirmation, FCO and AK) using the software provided by the OCT manufacturer (Eye Explorer 1.10.4 with viewing module 6.12; Heidelberg Engineering). Qualitative OCT data analyses were executed by or under supervision of a board-certified neuro-ophthalmologist (AJG).

#### Magnetic resonance tomography

Magnetic Resonance Imaging (MRI) data were available for 18 heterozygous *GRN* mutation carriers cross-sectionally and nine heterozygous *GRN* mutation carriers longitudinally with a follow-up of [median(min-max)] 1.4 (0.9-4.0) years. Participants were scanned on either a 3 Tesla Siemens Prisma fit or TrioTim at the UCSF Neuroscience Imaging Center. Magnetization Prepared Rapid Gradient Echo (MP-RAGE) sagittal images were acquired using the following parameters: 240 × 256 × 256 matrix; 160 slices; voxel size = 1.0 × 1.0 × 1.0 $mm^3$; flip angle = 9°; TR = 2300 ms; TE = 2.9 ms (Prisma) and TE = 2.98 (Trio). Participants were scanned on the same scanner for all time points. The Fluid Attenuated Inversion Recovery (FLAIR) images were acquired using the following parameters: 256 x 256 matrix; 176 slices (Prisma) and 160 slices (Trio); in-plane resolution = 1.0 x 1.0 mm (Prisma) and in-plane resolution = 0.98 x 0.98 mm (Trio); flip angle = 120°; TR = 5000ms (Prisma) and TR = 6000ms (Trio); TE = 397ms (Prisma) and TE = 388ms; TI = 1800ms (Prisma) and TI = 2100ms (Trio). White matter hyperintensities were quantified on FLAIR-imaging using the Fazekas scale with a Fazekas score of 3 representing severe white matter disease – the low sample size for a Fazekas score of 3 limits the generalizability of this exploratory analysis [12].

#### Volumetric processing

Prior to pre-processing, all T1-weighted images were visually inspected for quality control. Images with excessive motion or image artifact were excluded. Tissue segmentation was performed using SPM12 (Wellcome Trust Center for Neuroimaging, London, UK, http://www.Fil.ion.ucl.ac.uk/spm) unified segmentation [13]. An intra-subject template was created by non-linear diffeomorphic and rigid-body registration proposed by the symmetric diffeomorphic registration for longitudinal MRI framework [14]. The intra-subject template was segmented also using SPM12's unified segmentation. A within-subject modulation was applied by multiplying the timepoints' jacobian with the intra-subject averaged tissues

[15]. A customized group template was generated from the within-subject average gray and white matter tissues and cerebrospinal fluid by non-linear registration template generation using Large Deformation Diffeomorphic Metric Mapping framework [13]. Modulated intra-subject gray and white matter were geometrically normalized to the group template and then smoothed 8mm full width half maximum using Gaussian kernel. Every step of the transformation was carefully inspected from the native space to the group template. For statistical purposes, linear and non-linear transformations between the group template space and International Consortium of Brain Mapping (ICBM) [16] were applied. Quantification of volumes in specific brain regions was accomplished by transforming a standard parcellation atlas into ICBM space and summing all modulated gray matter within each parcellated region of interest (ROI) [17]. Total intracranial volume (TIV) was calculated for each subject as the sum of the gray matter, white matter, and cerebrospinal fluid segmentation.

**Analysis of clinical data**

Statistical analyses were performed with R 3.6.1. Figures were created using R and Adobe Illustrator. For all calculations, statistical significance was established at $p < 0.05$. Group differences were tested with $x^2$ for sex and Wilcoxon-Mann-Whitney U test for age. Cross-sectional and longitudinal group differences in OCT and cognition were evaluated by t-test and linear mixed effect models accounting for within-subject inter-eye dependencies as a random effect (for OCT parameters) as well as for age and sex in non-matched subsets respectively. OPL/ONL-ratio (OPL divided by ONL) was not set as an outcome parameter a priori but included as a combined outcome parameter with potential future as prognostic biomarker. Composite scores were calculated as the average of sample-based z-scores for the following domains: memory (CVLT delay, Benson delay), executive functions (Trails, design fluency, digits backwards, lexical fluency, Stroop), visuospatial function (Benson copy, VOSP) and language (BNT, semantic fluency). Group differences in MRI parameters were assessed using unpaired t-test.

Comparisons of cognitive and regional atrophy data were corrected for multiple testing by Bonferroni-Holm method, if necessary.

**Mice**

Wildtype and *Grn*$^{-/-}$ mice on C57BL/6J background [18] were raised under 12-h cyclic light on a standard diet. In some studies, the C3aR inhibitor SB290157 was administered intraperitoneally (0.5 mg/kg) to 1-month-old mice 3 times/week for 8 weeks. Depending on the experimental paradigm, mice of various ages were euthanized using $CO_2$ asphyxiation, eyes were removed, and eyecups were processed for immunohistochemistry, live imaging, immunoblotting, or transcriptomics as previously detailed [19-22]. All studies were approved by University of California, San Francisco animal care and use authorities and in accordance to ARRIVE guidelines.

**Immunostaining of mouse retinal cryosections**

Mouse eyes were collected immediately after euthanasia, punctured at the ora serrata, and fixed in 4% PFA for 1 h at room temperature. Anterior portion of the eyes including the lens were then removed, and the eyecups incubated in 30% sucrose in PBS for 16 h at 4°C prior to embedding in OCT. For better visualization of microglia migration, cryosections of 30 μm thickness were used. Cryosections were blocked in 4% BSA for 2.5 h at room temperature, immunostained with rabbit anti-CD68 (Abcam, 1:100) for 16 h at 4°C, followed by AlexaFluor secondary antibody (1:500) for 2 h at room temperature. Retinal layers were distinguished via DAPI (1:200, 15 min at room temperature) and WGA staining (10 μg/ml for 10 min followed by AlexaFluor488-conjugated streptavidin for 30 min at room temperature). Images were acquired on Nikon spinning disk confocal microscope with 10x (0.45 NA) or 20x (0.75 NA) objectives. For outer nuclear layer thickness analysis, mid-peripheral region of the retina was assessed, and the layer thickness measured using the "Distance Measurement" tool in Nikon NIS-Elements software.

**Statistics**

For mouse studies, data were analyzed by unpaired two-tailed *t*-test with Welch's correction for two experimental groups, and one-or two-way ANOVA tests with post-hoc analysis for comparison between multiple groups (Graphpad Prism). Data are presented as Mean ± SEM, with ≥ 3 mice per age, genotype, or treatment.

**Table S6. Key resources**

| REAGENT or RESOURCE | SOURCE | IDENTIFIER | RRID |
|---|---|---|---|
| Antibodies | | | |
| Actin | Santa Cruz Biotechnology Inc., Dallas, Texas | Cat. No. Sc-1616 | AB_2630402 |
| CD68 | Abcam, Cambridge, MA | Cat. No. ab125212 | AB_10975465 |
| Iba1 | Abcam, Cambridge, MA | Cat. No. ab5076 | AB_91676 |
| Iba1 | ThermoFisher, Waltham, MA | Cat. No. PA5-27436 | AB_2544912 |
| Rhodopsin N-terminus (clone 4D2) | Millipore, Burlington, MA | Cat. No. MABN15 | AB_10807045 |
| Rhodopsin C-terminus (clone 1D4) | Millipore, Burlington, MA | Cat. No. MAB5356 | AB_2178961 |
| Cathepsin D | R&D systems, Minneapolis | Cat. No. MAB1029 | AB_2292411 |
| LAMP1 | Sigma-Aldrich, St. Louis, MO | Cat. No. L1418 | AB_477157 |
| Oma1 (H-11) | Santa Cruz Biotechnology Inc., Dallas, Texas | Cat. No. SC-515788 | AB_2905488 |
| p-mTOR (Ser2448) | Cell Signaling, Danvers, MA | Cat. No. 5536 | AB_10691552 |
| 4E-BP1 (phospho-Thr37/46) | Cell Signaling, Danvers, MA | Cat. No. 2855 | AB_560835 |
| 4E-BP1(53H11) | Cell Signaling, Danvers, MA | Cat#9644 | AB_2097841 |
| MTP18 | Abcam, Cambridge, MA | Cat. No. ab198217 | N/A |
| C3 | ThermoFisher, Waltham, MA | Cat. No. ICN55444 | AB_2334469 |
| C3a | Abcam, Cambridge, MA | Cat. No. ab11873 | AB_298655 |
| C3aR | Novus Biologicals, CO | Cat. No. NBP2-15649 | N/A |
| TOM20 | Santa Cruz Biotechnology Inc., Dallas, Texas | Cat. No. sc-11415 | AB_2207533 |
| OPA-1 | Novus Biologicals, CO | Cat. No. NB110-55290 | AB_829789 |
| Mfn2 | Abcam, Cambridge, MA | Cat. No. ab56889 | AB_2142629 |
| Phospho-DRP1 (Ser637) | Cell Signaling, Danvers, MA | Cat. No. 4867S | N/A |
| PhosphoDRP1 (Ser616) | ThermoFisher, Waltham, MA | Cat. No. PIPA564821 | N/A |
| FIS1 | Proteintech, Rosemont, IL | Cat. No. 10956-1-AP | AB_2102532 |
| RPE65 | Abcam, Cambridge, MA | Cat. No. ab13826 | AB_2181006 |
| Chemicals, peptides, and recombinant proteins | | | |
| NativePAGE™ 4-16% Bis-Tris gel | Invitrogen, Waltham, MA | Cat. No. BN1004BOX | |

| | | |
|---|---|---|
| NativePAGE™ Sample Prep Kit | Invitrogen, Waltham, MA | Cat. No. BN2008 |
| NativePAGE™ Running Buffer Kit | Invitrogen, Waltham, MA | Cat. No. BN2007 |
| NativeMark™ Unstained Protein Standard | Invitrogen, Waltham, MA | Cat. No. LC0725 |
| NuPAGE 4-12% Bis-Tris | Invitrogen, Waltham, MA | Cat. No. NP0336BOX |
| 4X NuPAGE LDS sample buffer | Invitrogen, Waltham, MA | Cat. No. NP0007 |
| 10X NuPAGE Sample Reducing Agent | Invitrogen, Waltham, MA | Cat. No. NP0009 |
| Phenylmethylsulfonyl fluoride (PMSF) | Tocris, Bristol, UK | Cat. No. 4486 |
| Protease Inhibitor Cocktail Set III, EDTA-Free | Millipore, Burlington, MA | Cat. No. 539134 |
| Intercept (PBS) Protein-Free Blocking Buffer | LI-COR, Lincoln, NE | Cat. No. 927-90001 |
| Revert™ 700 Total Protein Stain and Wash Solution Kit | LI-COR, Lincoln, NE | Cat. No. 926-11015 |
| SeeBlue Plus2 Pre-stained Protein Standard | Invitrogen, Waltham, MA | Cat. No. LC5925 |
| Bovine Serum Albumin | Rockland Immunochemicals, Pottstown, PA | Cat. No. BSA-50 |
| Calcium Chloride Dihydrate | Sigma-Aldrich, St. Louis, MO | Cat#C7902 |
| DAPI (14.3 mM stock solution, used at 1:200) | Sigma-Aldrich, St. Louis, MO | Cat. No. D9542 |
| Glucose | Sigma Aldrich, St. Louis, MO | Cat. No. G7528 |
| HBSS | Corning, Corning, NY | Cat. No. 21-023-CV |
| HEPES | ThermoFisher, Waltham, MA | Cat. No. 15630080 |
| Laemmli SDS-Sample Buffer | BioWorld, Dublin, OH | Cat. No. C995V98 |
| Magnesium Chloride Hexahydrate | Sigma-Aldrich, St. Louis, MO | Cat. No. M2393 |
| Paraformaldehyde (8%) | Electron Microscopy Sciences, Hatfield, PA | Cat. No. 157-8 |
| Phosphate Buffer Saline | ThermoFisher Scientific, Waltham, MA | Cat. No. BP665-1 |
| 10X RIPA buffer | Abcam, Cambridge, MA | Cat. No. ab156034 |
| Saponin | Sigma-Aldrich, St. Louis, MO | Cat. No. 84510 |
| Sucrose | Sigma-Aldrich, St. Louis, MO | Cat. No. S0389 |
| TrueBlack | Biotium, Fremont, CA | Cat. No. 23007 |
| VectaShield | Vector Laboratories, Burlingame, CA | Cat. No. H1000 |
| Ciprofloxacin | Sigma-Aldrich, St. Louis, MO | Cat. No. 17850-5G-F |

| | | |
|---|---|---|
| DMEM | Corning, Corning, NY | Cat. No. 10-013-CV |
| Fetal Bovine Serum (heat-inactivated) | American Type Culture Collection, VA | Cat. No. 30-2020 |
| Non-essential amino acids (NEAA) | Corning, Corning, NY | Cat. No. 25-025-CI |
| Penicillin-Streptomycin | Corning, Corning, NY | Cat. No. 30-002-CI |
| 0.25% Trypsin | Corning, Corning, NY | Cat. No. 25-053-CI |
| 2.5% Trypsin | Lonza, Walkersville, MD | Cat. No. 17-160E |
| Opti-MEM | Gibco, ThermoFisher, Waltham, MA | Cat. No. 31985-070 |
| SB290157 | Tocris Bioscience | Cat. No. 6860 |
| Critical commercial assays | | |
| DC protein assay reagents | Bio-Rad, Hercules, CA | Cat. No. 5000116 |
| Experimental models: Organisms/strains | | |
| C57BL/6J mice | Jackson labs | Strain #: 000664 RRID:IMSR_JAX:000664 |
| *Grn*$^{-/-}$ mice (B6.129S4(FVB)-*Grn*$^{tm1.1Far}$/Mmjax) | UCSF Gladstone Institute | Strain #: 036771 RRID: B6.129S4(FVB)-*Grn*$^{tm1.1Far}$/Mmjax |
| Software and algorithms | | |
| GraphPad Prism | GraphPad Prism®, La Jolla, CA | |
| R (Versions 3.6.2–4.1.1) | The R Foundation | https://www.r-project.org/foundation/; RRID:SCR_001905R |
| Adobe Illustrator | Adobe, San Jose, CA | |
| Photoshop | Adobe, San Jose, CA | |
| Imaris | Bitplane, Concord, MA | |
| Microsoft Excel | Microsoft Corporation, Albuquerque, NM | |
| NIS Elements | Nikon, Melville, NY | |

| Dyes | Concentration | Source | Catalog number |
|---|---|---|---|
| DAPI | 14.3 mM stock solution, used at 1:200 | Sigma-Aldrich, St. Louis, MO | D9542 |
| Acti-stain 488 phalloidin | 1:200, incubated with secondary antibodies | Cytoskeleton Inc., Denver, CO | PHDG1 |

| | | | |
|---|---|---|---|
| Mitotracker Deep Red | 200 nM, 15 min | ThermoFisher, Waltham, MA | M22426 |
| BioTracker ATP Red | 1:1000, 15 min | Sigma-Aldrich, St. Louis, MO | SCT045 |
| TMRE | 500 nM, 10 min | Biotium, Fremont, CA | 70005 |
